# Distinct neural dynamics in prefrontal and premotor cortex during flexible perceptual decisions

**DOI:** 10.64898/2026.02.02.702013

**Authors:** Tian Wang, Eric Kenji Lee, Nicole Carr, Yuke Li, Chandramouli Chandrasekaran

## Abstract

The neural mechanisms underlying flexible perceptual decisions, where the mapping between sensory input and motor action changes depending on context, remain unclear. Here we show that prefrontal and premotor cortex use distinct neural dynamics to implement different computations during flexible decisions. We trained monkeys to discriminate the dominant color of a red-green checkerboard and report their decision by choosing one of two targets^1;2^. By randomizing the target configuration on a trial-by-trial basis, we ensured a flexible mapping between color (red vs. green) and action choice (left vs. right), necessitating a nonlinear exclusive- or (XOR) computation^3–5^. We found that neural dynamics in dorsolateral prefrontal cortex (DLPFC) led to higher-dimensional population representations than those in dorsal premotor cortex (PMd). Neural activity in DLPFC first separated by target configuration, then by color choice and action choice after stimulus onset, reflecting the XOR computation. In contrast, neural dynamics in PMd led to lower-dimensional representations that only reflected action choice, the output of the XOR computation. These higher-dimensional representations in DLPFC enabled earlier decoding of both color choice and action choice compared to PMd, and were strongest in anterior and ventral DLPFC^6^. These findings reveal distinct computations by neural dynamics: prefrontal cortex implements flexible sensorimotor mappings through high-dimensional representations while dorsal premotor cortex reflects only the selected action.

---

We have an excellent understanding of the decision-related neural computations in tasks with a fixed mapping between stimulus and response^7–11^. In contrast, we currently do not understand how flexible perceptual decisions emerge in the brain when the mapping between sensory stimulus and motor response changes depending on context^3;5;12^. For example, depending on the intersection you are at (the context), a green signal might mean turning left or turning right to get home. Recent studies have shown that perceptual decision signals (whether the light is red or green) can be dissociated from motor responses (turn left vs. right) when stimulus-response mappings are flexible^3;4^. Yet, how neural circuits perform relevant computations for such flexible decisions, and whether this computation is more localized to some brain areas or distributed across many areas, remains unresolved^12;13^.

First, what computations do neural circuits perform to enable flexible decisions? When the same sensory stimulus necessitates different actions based on context, neural circuits must solve an XOR (exclusive-or) computation (Fig. 1a)^5^. One potential solution is for neural dynamics in a brain area to create time-varying higher-dimensional representations encoding conjunctions of the perceptual decision and the potential action choice, allowing the correct action to be linearly decoded for the different contexts. Therefore, a key signature of a brain area that implements this computation is that it must encode context, perceptual decision, and action choice. Whether neural dynamics in relevant brain areas have such signatures is currently not resolved.

**Figure 1:**
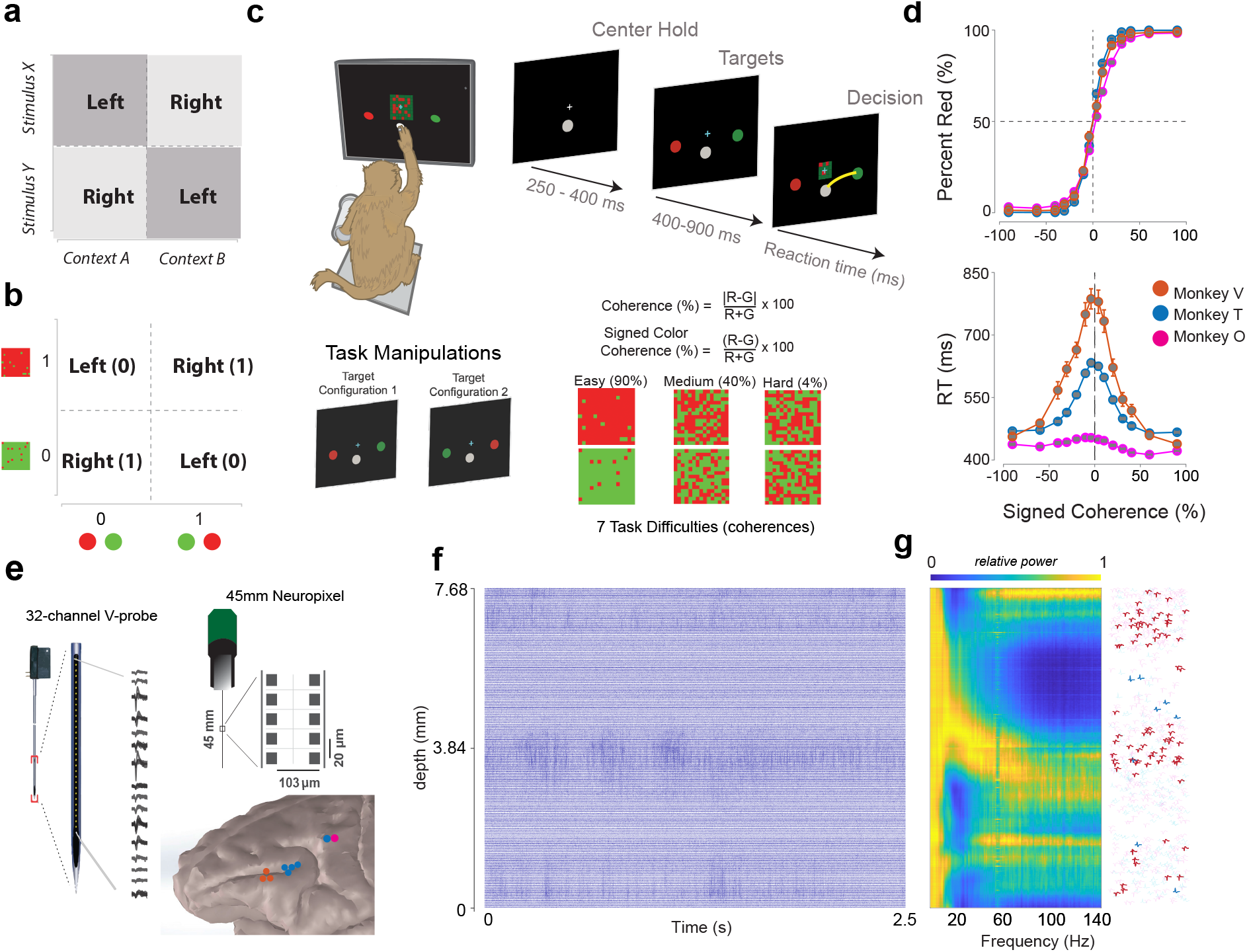
Monkeys can flexibly respond to sensory stimuli based on context: (**a**) Schematic illustrating that the flexible mapping between sensory input (stimulus X or Y) and action choice (left or right) based on context (context A or B) needs an XOR computation of sensory input and context. (**b**) Action choice in *our task* emerges from an XOR computation on color (red vs. green) and target configuration (**red-left&green-right**/**green-left&red-right**). (**c**) An illustration of the behavioral task setup (top left), timeline of the task (top right) and task manipulations (bottom). Top left, setup: Monkeys sat comfortably in the primate chair to perform the task and the middle finger of their active hand was taped with an infrared light reflective bead to track the hand position in 3 dimensions. Top right: Timeline of the target first checkerboard discrimination task. Bottom: Two task manipulations. For each trial, we used one of the two target configurations randomly, and a random checkerboard cue composed of a mixture of red and green squares. The checkerboard was parameterized by color coherence, defined as 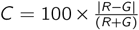. The corresponding signed coherence (*SC*) is defined as 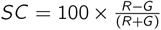. Positive values of signed coherence (*SC*) denote more red than green (G) squares and vice versa. (**d**) Psychometric curves, percent responded red (top), and reaction time (RTs for all trials, bottom) as a function of the signed coherence of the checkerboard cue, averaged over sessions for three monkeys (see 2) Top: psychometric curves of monkey T, V, and O. Bottom: RT curves for the same recording sessions. Error bars represent SEM. (**e**) Left: Schematic of 32-channel Plexon V-probe. Right top: Schematic of imec Neuropixels 1.0 NHP. Right bottom: recording sites from DLPFC and PMd of 3 monkeys (monkey T, V, O) localized using T1 MRI scans. (**f**) A 2.5 s snippet of the spike band (300 Hz - 6 KHz) of 384 channels covering 3 cortices (putatively Brodmann area 8Ad, 9/46d and 9/46v) in a single recording session. (**g**) Left: spectrolaminar analysis of local field potentials enables delineation of multiple brain areas in DLPFC. Right: location of all spike clusters as a function of depth. The opacity denotes clusters modulated robustly in the task.

Second, are these computations distributed across these brain areas, or are some regions specialized for the XOR computation needed for combining context and sensory evidence while others represent motor actions?^13^. Studies using fixed stimulus-response mappings suggest decision-variables are found in many brain regions including premotor brain areas^1;2;14–17^. In contrast, human studies suggest a functional gradient with prefrontal cortex supporting abstract computations and premotor cortex representing concrete action plans^18^. Similarly, visual encoding also varies across areas of the prefrontal cortex of monkeys^19^. Whether similar specialization exists for flexible decisions is unclear.

To address these questions, we took inspiration from past studies of flexible decisions^3;4;20–22^ and trained monkeys in a task that varied the mapping between sensory input and motor action on a trial-by-trial basis. In this task, monkeys discriminated the dominant color in a central static checkerboard composed of red and green squares and reported their decision by reaching to and touching a target of the corresponding color. By randomizing the target configuration, we created two contexts that reverse the mapping between the color of the checkerboard and action choice: in one context, red means reach left and green means reach right, while in the other context, red means reach right and green means reach left. Identifying the correct action choice in this task again requires a nonlinear XOR computation of the sensory input (the color) and the context (target configuration) (Fig. 1b, c).

If a brain region is performing computations for flexible decisions, we would expect three hallmarks in its neural activity. First, the brain area should encode all task variables (context, color, and action) for flexible mapping between stimulus and response, thus enabling high-dimensional representations that can unmix perceptual decisions from action. Second, color choice should precede action choice, reflecting the computation of combining color and context to identify action choice. Third, and finally, behaviorally-relevant representations should emerge earlier in this brain area than in downstream motor areas, which only represent the outputs of the computations. We found that DLPFC but not PMd exhibited all three of these hallmarks^1;17;23–25^. DLPFC neurons encoded context, color, and action, and the interaction between color and action (also referred to as nonlinear mixed selectivity^26^). Color signals emerged before action signals, and finally action representations in DLPFC preceded those in PMd. These patterns emerged because neural dynamics in DLPFC led to timevarying high-dimensional representations that covaried with both color choice and action choice, consistent with implementing the XOR computation^6;26;27^. In contrast, PMd dynamics reflected only action choice. Selectivity for color and interaction between color and action were strongest in ventral and anterior DLPFC. These findings reveal a functional gradient from abstract computations in prefrontal cortex to action-related dynamics in premotor cortex.

## Results

### Monkeys can perform flexible decisions

We trained three monkeys (T, V, and O) to perform a flexible reaction time decision-making task that decouples perceptual decisions from action selection (Fig. 1a-b). Monkeys discriminated the dominant color in a central red-green checkerboard and reported their decision with an arm movement (Fig. 1c, top panel). We used two task manipulations inspired by past studies^3;4;20;21^. First, we randomized the target configuration (color of left and right targets, Fig. 1c, bottom left panel) on a trial-by-trial basis, creating a flexible mapping between sensory input (checkerboard color) and action (reach direction) that necessitates an XOR computation. This manipulation enabled rigorous dissociation of how neural activity varies as a function of both color inputs/choice (red vs. green) vs. the output of the computation, the action choice (left vs. right). Such dissociation is not possible with fixed stimulus-response mappings. Second, we varied discrimination difficulty (Fig. 1c, right panel), inducing errors and producing a range of reaction times and bounding the time window for decision-related dynamics. Despite this added complexity, monkeys performed the task well, reporting red for red-dominant checkerboards and green for green-dominant checkerboards, with reation time (RTs) that varied logically with sensory evidence (Fig. 1d).

While animals performed the task, we recorded single neurons and multi-units from various aspects of the prefrontal cortex along the principal sulcus (monkeys T, V) and PMd (T, O) using single electrodes, 32-channel multi-contact electrodes and 10/45 mm NHP 1.0 Neuropixels (Fig. 1e). We recorded from the superficial aspects of DLPFC (areas 8Ad) and deep inside the principal sulcus (9/46d or DLPFCd, and 9/46v or DLPFCv) typically during different sessions (Fig. 1e, right bottom panel). We localized the recording locations using structral MRI, spectrolaminar analysis, and cortical depth (Fig. S1a-d)^28^. In a few sessions, we also used the 45 mm Neuropixels to record from multiple DLPFC areas simultaneously in a single session (Fig. 1f-g). PMd recordings were mostly restricted to caudal aspects of PMd areas (PMdc). In total, the DLPFC dataset reported here included 5570 single neurons and multi-units (4863 from T, 707 from V), and our PMd dataset consisted of 1686 units (1236 from T, 450 from O). To gain additional power for some of the analyses of how encoding varies as a function of the subregions of the brain (Fig. 5), we also included data from a fourth monkey Z (Fig. S1d, e). Data from monkey Z is not used for any other analyses.

### Distinct decision-related neural dynamics in DLPFC and PMd

DLPFC and PMd neurons differed in two key ways in this task. First, DLPFC neuronal responses were richer than those of PMd. Fig. 2 illustrates 4 example units from DLPFC (Fig. 2a-d) and 4 example units from PMd (Fig. 2e-h). DLPFC neurons modulated their firing rates to all the variables relevant for flexible decisions: target configuration, color choice (red vs. green, hereafter color), and the action choice (left vs. right, hereafter action). Target configuration-related firing rate changes in DLPFC neurons could be transient, sustained, or both (Fig. 2a-d). In stark contrast, PMd neurons modulated their responses primarily to action with minimal modulation to target configuration or color (Fig. 2e-g).

**Figure 2:**
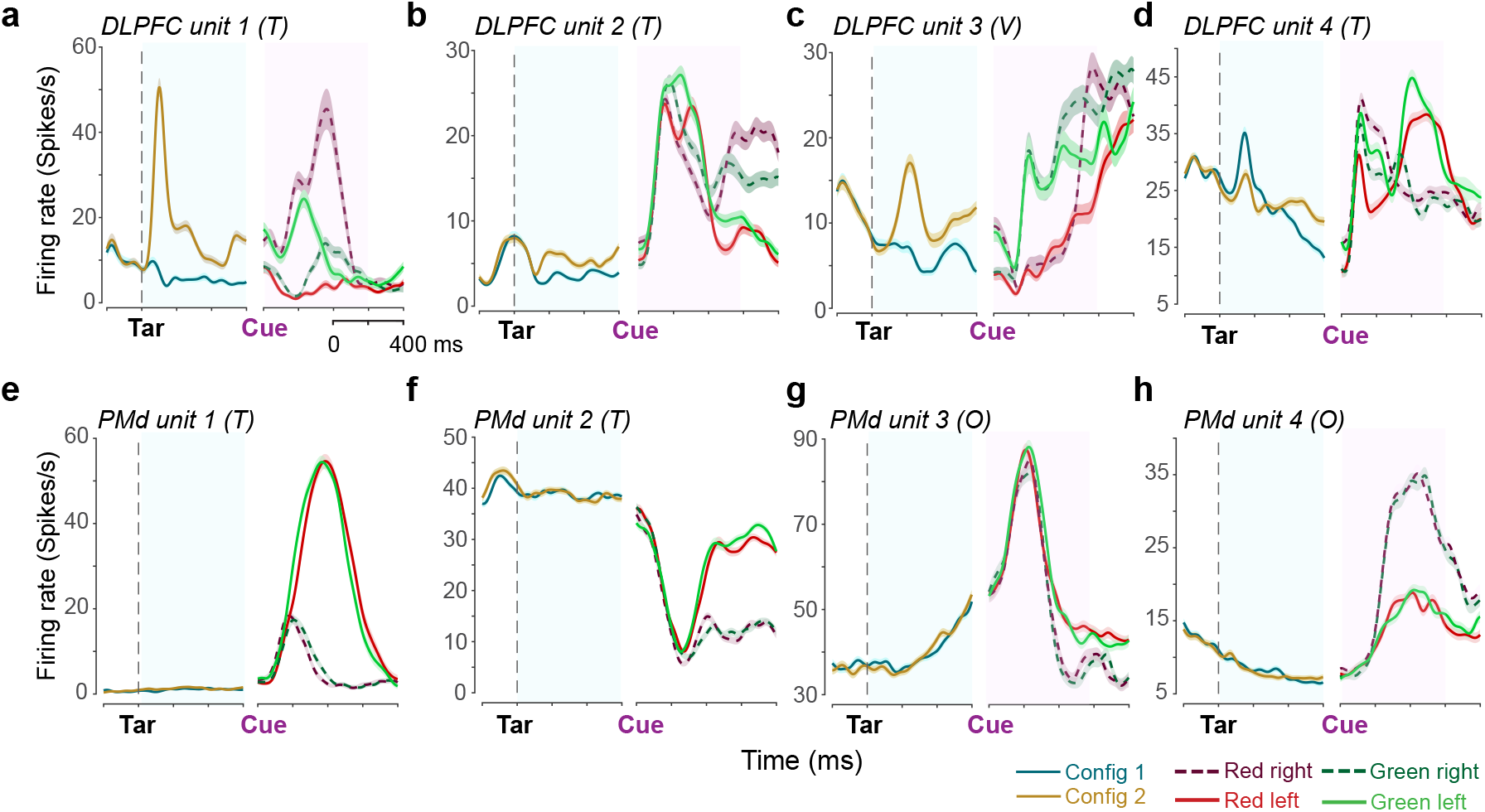
DLPFC neurons encodes task variables relevant for the XOR computation needed for flexible decisions. (**a-d**) Average PSTH for four example DLPFC units (labeled by area, unit ID, and monkey) showing responses aligned to target onset (Tar) and checkerboard onset (Cue). Solid lines denote left reaches, dashed lines denote right reaches (**red-left**(—), **red-right**(----), **green-left**(—), **green-right**(----)). DLPFC neurons modulate their firing rates as a function of the target configuration (or context, as in **a-d**), color choice (as in **c**), and action choice (as in **d**), and most-importantly separate by the combinations of color and action choice (i.e., mixed selectivity, as in **a-d**). (**e-h**) Same as (a-d) but for four example PMd units. Unlike DLPFC, PMd neurons primarily encode action choice (left vs. right) with minimal sensitivity to target configuration or color choice. Shaded areas represent SEM for all panels.

Second, DLPFC units were often modulated by more than one task variable (Fig. 2a,c,d), and most importantly showed sensitivity to combinations of color and action. Such tuning of DLPFC neural responses to combinations of variables is a hallmark of nonlinear mixed selectivity (NMS, see Appendix A: Nonlinear-Mixed Selectivity)^6;26;27;29^. Consequently, DLPFC neuronal responses separated as a function of both color and action, in many cases producing four distinct firing rate trajectories after checkerboard onset (Fig. 2a,b,d; see Fig. S2a-f for additional examples).

We summarized these single neuron responses using a two-way ANOVA with color and action as main effects, aligned to checkerboard onset, and estimated the percentage of significantly modulated units in DLPFC and PMd (see *Methods: single-unit encoding of task variables* and Appendix A: Nonlinear-Mixed Selectivity)^30^. We also quantified the effect size using a normalized modulation index for each task variable. These analyses revealed that a greater proportion of DLPFC units modulated firing rates as a function of color, target configuration, and critically the interaction between color and action (color × action, that is nonlinear mixed selective), and with larger effect sizes than PMd units (Fig. S2g-j).

Collectively, these single neuron analyses demonstrate that DLPFC neurons encode target configuration, color, action, and color × action (nonlinear mixed selectivity). The key advantage of encoding multiple variables and nonlinear mixed selectivity is that the population can form high-dimensional representations that enable solving the XOR computation needed to perform this flexible decision-making task^6;26^. We describe the properties of these high-dimensional representations in the next sections.

### Higher dimensional representations in DLPFC compared to PMd

The single neuron examples in Fig. 2 suggest greater complexity of firing rates in DLPFC compared to PMd both in terms of the task variables encoded and in the firing rate profiles as a function of time. We hypothesized that this heterogeneity in PSTHs would lead to higher dimensional representations in DLPFC^6;31^ compared to PMd, which, in turn, would enable robust solutions to the XOR problem. Consistent with this hypothesis, ∼6 principal component (PC) dimensions were required to explain 90% of the variance for *trial-averaged* Gaussian smoothed firing rates of DLPFC neurons, but only 3 dimensions were required for PMd neurons (Fig. 3a, top panel, p < 0.02, estimated from 100 bootstrap samples for *N*_*neurons*_ ≥ 100, see *Methods: Population dynamics*). This relationship held even after subtracting the across-condition average signal (Fig. 3a, bottom panel), indicating that this higher dimensionality in DLPFC emerges from differences in encoding of task variables.

**Figure 3:**
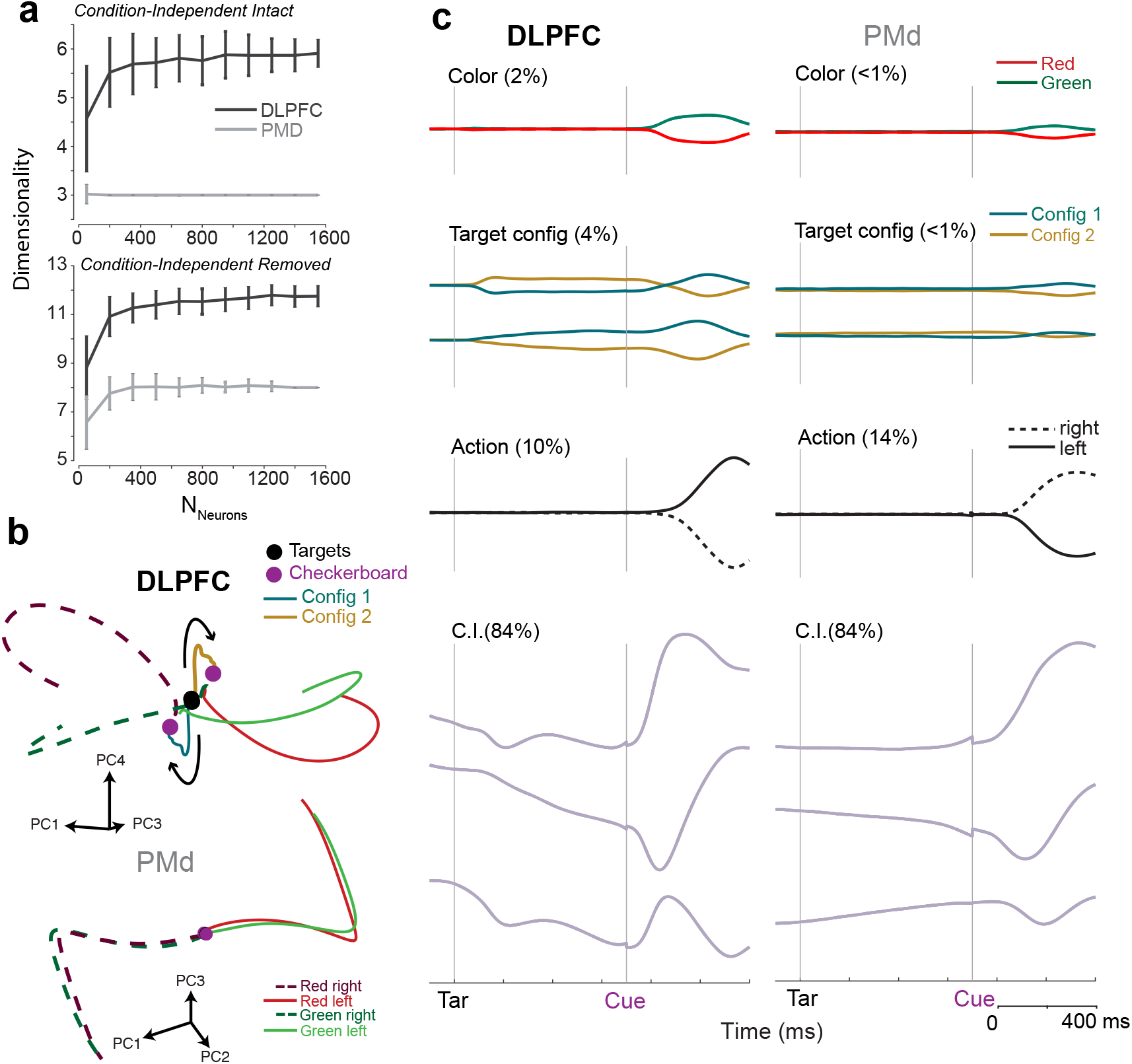
Neural population dynamics in DLPFC lead to higher dimensional representations that encode multiple task variables (a) Intrinsic dimensionality of population firing rates in DLPFC is larger than that of PMd. Number of principal components required to explain 90% of variance with respect to number of units sub-sampled from DLPFC and PMd dataset. Top: intact firing rates show DLPFC requires ∼6 dimensions while PMd requires ∼3. Bottom: after removing condition-independent (C.I.) signals by subtracting mean responses across the four task conditions, DLPFC still requires more dimensions (∼11) than PMd (∼8), indicating genuine multi-dimensional task encoding beyond temporal complexity. Error bars represent standard deviation from bootstrapping (n=100). (**b**) PCA trajectories for concatenated target and checkerboard epochs. In DLPFC (top), population activity separates into two trajectories corresponding to the two target configurations after target onset (black dot). Following checkerboard onset (magenta dot), each trajectory separates based on color choice, yielding four distinct trajectories (**red-left, red-right, green-left, green-right**). Trajectories ultimately converge based on action choice, reflecting the XOR computation required by the task. In PMd (bottom), population activity remains largely overlapping during target presentation and separates only by action choice after checkerboard onset. (**c**) Demixed principal component analysis (dPCA) reveals distinct task variable encoding in each area. Each row shows dPCA components for different task variables: color choice (top), target configuration (2nd row), action choice (3rd row), and condition-independent signals (bottom). Percentages indicate variance explained by each component. In DLPFC (left), color choice (2%), target configuration (4%), and action choice (9%) all explain substantial variance. In PMd (right), only action choice explains significant variance (13%), while color (<1%) and targe configuration (<1%) explain minimal variance. Vertical gray lines indicate target onset (Tar) and checkerboard onset (Cue).

The results from Fig. 3a predict greater amount of task-variable related signals in DLPFC compared to PMd during flexible decisions. We tested this prediction in two ways. We first visualized the PCs calculated from trial-averaged firing rates after subtracting condition-independent signals from each neuron. We found that target onset led to distinct neural dynamics in DLPFC states for each target configuration. The appearance of the checkerboard led to further evolution in time with two trajectories evolving from each target configuration (Fig. 3b, top panel). Thus, in DLPFC, neural activity separates by all possible behavioral outcomes (**Red Left, Red Right, Green Left, and Green Right**), which eventually converge together for action. In contrast, activity in PMd was largely unmodulated during the target viewing period and only separated as a function of action after checkerboard onset (Fig. 3b, bottom panel).

Second, we quantified task-variable related variance using demixed principal component analysis (dPCA,^32^). Variance for target configuration (4%), and color choice (2%) for DLPFC was significantly different from 0 (p < 0.02, permutation test) and exceeded the variance for these task variables in PMd (< 1%, p < 0.002, bootstrap test, n=1000, comparing dPCA variance for color and target configuration using sub-sampled DLPFC populations and PMd, see *Methods: Population dynamics*). In PMd, dPCA revealed that the only significant task variable that explained any variance was the action choice (Fig. 3c, top panel). Variance for action choice was higher in PMd than in DLPFC (p = 0.025, N=1000, bootstrap test comparing variance for action for sub-sampled DLPFC populations and PMd). Finally, condition independent signals (Fig. 3c, bottom panel) also showed a consistent difference between DLPFC and PMd. In DLPFC, condition independent signals changed rapidly both after target and checkerboard onset, but PMd activity only changed after checkerboard onset.

These results demonstrate that neural dynamics and representations in DLPFC are higher dimensional, covary with target configuration, and unmix color from action, whereas neural dynamics in PMd are lower dimensional and only covary with action.

### Neural dynamics in DLPFC leads to representations that solve the XOR problem

Our analyses in Fig. 3 suggest higher dimensional representations encoding multiple task variables in DLPFC. We next investigated the geometry of these representations and how they changed with time, on both correct and error trials. We first visualized the low-dimensional neural activity of 4 color/action combinations at different time points around checkerboard onset (Fig. 4a).

**Figure 4:**
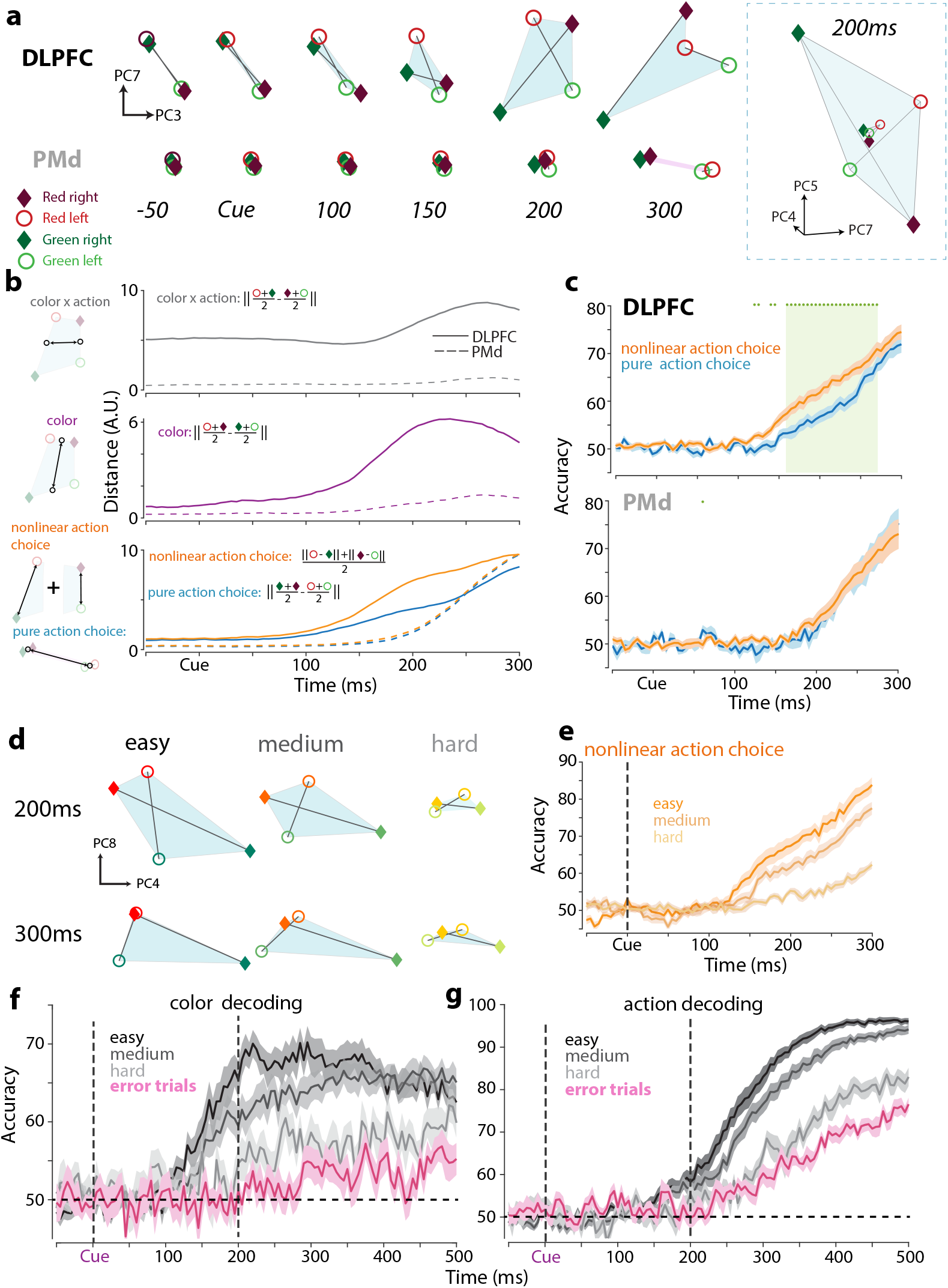
Neural representations in DLPFC separate by color and action. (**a**) Low-dimensional visualization of population dynamics. Projections onto PC3 and PC7 for DLPFC (top) and PMd (bottom) relative to checkerboard cue onset. In DLPFC, the four conditions (red-left, red-right, green-left, green-right) are well separated and form a quadrilateral in 2 dimensions. In PMd, only action choice (filled vs. open circles) separates the population state, with minimal color-related structure, as opposed to DLPFC, where color also becomes separable. Right panel shows a 3D visualization of the DLPFC (large tetrahedron) and the PMd (small tetrahedron) state at 200 ms. **(b)** Euclidean distance in the top 10 PCs (>90% variance explained) for interaction term of color × action, color, nonlinear action choice, and pure action choice. DLPFC (solid line) shows substantial and sustained separation for the color × choice interaction (gray) and color (purple), while PMd (dashed line) shows minimal separation. Both areas show increased separation for choice-related signals (orange: nonlinear choice; blue: pure choice), with PMd showing only pure choice separation. (**c**) Average logistic regression decoding accuracy for nonlinear choice (orange) and pure choice (blue) using only Neuropixels sessions (n=23 sessions DLPFC, n=12 sessions in PMd). Shaded region indicates significant differences between nonlinear choice and pure choice as assessed by a paired t-test for each bin (p < 0.05). (**d**) PC projections (PC4 and PC8) at 200 ms (top) and 300 ms (bottom) for easy (high coherence), medium, and hard (low coherence) trials. Note the separation between potential color and action combinations is stronger for easier compared to harder trials. (**e**) Nonlinear choice decoding accuracy decreases with stimulus difficulty. Shaded regions indicate SEM across sessions for DLPFC. (**f**) Color decoding accuracy across coherence levels. Easy trials show sustained high accuracy (∼70%) while medium and hard trials show lower accuracy, with error trials (magenta) performing near chance. Vertical dashed line at 200 ms is provided to compare (f) and (g). (**g**) Action choice decoding accuracy across coherence levels. All correct trial types reach high accuracy (>90%) while error trials (magenta) show chance-level performance until later in the trial when the monkey ultimately makes an (incorrect) choice. Vertical dashed line at 200 ms is provided to compare (f) and (g).

The first two principal components were dominated by condition-independent signals, but higher dimensions starting from PC3 onwards encoded variables relevant for flexible decisions. Fig. 4a shows the trial-averaged firing rates projected onto PC3 and PC7 (Fig. S3 visualizes these conditions in PC4 and PC5). Before checkerboard onset, population activity in DLPFC was modulated by the two target configurations (**RL&GR** separated from **RR&GL**). After checkerboard onset (∼150 ms), color-related signals emerged in DLPFC. The low-dimensional neural representation of 4 color/action combinations are separated from each other (**red-left, red-right, green-left**, and **green-right**), forming a quadrilateral in two dimensions and a tetrahedron in three and higher dimensions, in line with nonlinear mixed selectivity in single neurons^26^. Both the volume and surface area of the tetrahedron formed by the four color-action combinations in the top 10 PCs (capturing *>* 90% of variance) increased in DLPFC after checkerboard onset (Fig. S3d-e), reflecting increasing separation of all four conditions^6;26;31^.

In contrast, neural representations of task variables in PMd showed only simple geometric structure (bootstrap test comparing volume and surface area of subsampled DLPFC dataset vs. PMd, p < 0.02, n=100 bootstrap samples). Before target onset, all four conditions were intermixed, and this pattern continued even after checkerboard onset. Only ∼200 ms post-checkerboard did activity begin to separate based on pure action signals. These visualizations confirm high-dimensional representations in DLPFC that unmix color and action, whereas PMd primarily encodes action.

We quantified this geometric structure by measuring Euclidean distances between conditions in the PC space that explained more than 90% of the variance. The distance between color × action interaction terms (e.g., **RL&GR** vs. **RR&GL**) strongly increased in DLPFC after checkerboard onset (Fig. 4b), but only weakly in PMd. A similar pattern emerged for color distance (R vs.G) (bootstrap test comparing subsampled DLPFC dataset and PMd, p < 0.02 for both color and color × action, n = 100 bootstrap samples). We defined “nonlinear choice” as the average separation between **RL** vs. **GR** and **RR** vs. **GL**, and “pure (linear) choice” as the separation between left vs. right. If mixed selectivity is present, then all 4 color/choice combinations are separated from each other: nonlinear choice should exceed pure choice. In contrast, if the population only has pure selectivity, then nonlinear choice and pure choice should be similar (see Fig. S4a for a detailed explanation). In DLPFC, nonlinear choice was larger than pure choice, whereas in PMd, the two were nearly identical (Fig. 4b, bottom panel, bootstrap test using a subsampled DLPFC population to compare nonlinear choice and pure choice in the 150-250 ms period after checkerboard onset, p < 0.02, N = 100 bootstrap samples).

Decoding analyses further confirmed that color and action were separable at the single-trial level. In DLPFC, we could decode nonlinear choice earlier (nonlinear choice: 120 ± 11 ms, pure action choice: 175 ± 14 ms, permutation test, p < 0.0015, Fig. 4c) and stronger than pure action choice (paired t-test, p < 0.05, see green shading). Around 300 ms after checkerboard onset, these signals began to converge (as in Fig. 4a, DLPFC 300 ms). In contrast, PMd showed nearly identical and *delayed* time courses for both decoders (nonlinear choice: 200 ± 14 ms, pure action choice: 205 ± 10 ms, p = 0.33, permutation test), consistent with only pure action-related signals (Fig. S4a). These choice signals also emerged later in PMd compared to DLPFC (bootstrap test, n=10000 bootstrap samples, comparing latency for mixed and pure choice selectivity for DLPFC compared to PMd pure choice latency, p <.002 for mixed, and p <.006 for pure). The presence of robust mixed selectivity in DLPFC also predicted that all pairwise condition combinations should be decodable with similar timecourses (Fig. S4b). Indeed, after checkerboard onset, decoding accuracy largely followed a similar time course for all pairwise combinations in DLPFC but not in PMd (Fig. S4c).

These high-dimensional representations also changed with stimulus difficulty and on error trials. First, the area bounded by the four conditions decreased with increasing difficulty (Fig. 4d). This pattern was also observed in other PC dimensions (Fig. S3b) and when aligned to movement onset (Fig. S3c). At the single-trial level, nonlinear choice decoding accuracy decreased significantly as task difficulty increased (Fig. 4e, one-way ANOVA on average decoding accuracy between 100 and 300 ms, F(2, 66) = 39.92, p=8.6×10^−12^), consistent with weaker mixed representations for harder trials.

These decoding analyses also revealed two other features in the data: First, we could decode color shortly after checkerboard onset (140 ± 15 ms) and the accuracy of this decoding depended on the difficulty of the checkerboard (one-way ANOVA on average decoding accuracy between 100 and 300 ms, F(2, 45) = 12.48, p=3.4×10^−5^). In addition, these color signals emerged earlier than action signals (p=0.021, permutation test, N=10000) and were stronger than action signals for the easiest and medium task difficulties in a 100 to 200 ms period after checkerboard onset — a period where the nonlinear choice signals are strong (paired t-tests, easy: t_22_=3.93, p < 0.0007, medium: t_22_=2.81, p < 0.01, hard: t_22_=1.66, p=0.10).

Second, neural representations on correct and error trials were different. We trained a color decoder on correct trials and tested on held-out correct and error trials for hard (low coherence) trials. Color decoding for error trials dropped to near chance levels and was lower for an equivalent set of correct trials in the 200 to 500 ms period after checkerboard onset (Fig. 4f, *t*_15_ = 3.60, p < 0.003). Consequently, action signals emerge slower for error trials than correct trials after checkerboard onset (Comparing average choice accuracy in the 200-500 ms after checkerboard onset Fig. 4g, *t*_15_ = 3.72, p < 0.002), suggesting that errors arise from weaker color representations in DLPFC.

### Color and mixed selectivity is stronger in anterior and ventral DLPFC

Our results so far demonstrate strong differences in dynamics between DLPFC and PMd during the TF task. We also investigated whether there is similar task modulation everywhere in DLPFC, or whether color and nonlinear mixed selectivity varies within different sub areas of DLPFC as predicted by anatomical and functional studies^18;19;33;34^. Past studies of DLPFC typically recorded from only a few hundred neurons, making such gradients difficult to detect. Our large dataset with > 7000 units (Table 2) across DLPFC and PMd enabled us to re-examine this question. We combined MRI and histology to estimate recording locations across DLPFC. In posterior recordings, we sampled from 8Ad (superficial) and 9/46d and 9/46v (deep) (Fig. S1a-d). In anterior recordings primarily from monkey V, we recorded from both banks of the principal sulcus. For each unit, we estimated its anterior-posterior position relative to posterior end of principal sulcus and its depth from dura. To add additional power for just these analyses, we also included data from a fourth monkey (monkey Z) where we could perform a few recordings in areas in and around the posterior aspect of the principal sulcus.

Fig. 5a-c shows a scatter plot of the absolute value of dPCA loading for each unit (obtained from dPCA on normalized firing rates) as a function of the anterior-posterior and depth location. The size of the dot represents the amplitude of each unit’s dPC loading on color, action, and color × action (nonlinear mixed selectivity; see Fig. S5a-c for the same dPCA loadings without normalization of firing rates or using effect size (Fig. S5d-f). In Fig. S5h top, we also plotted each unit’s mixed selectivity index to quantify the joint representation strength of color, action and color × action. Color and mixed selectivity were strongest in anterior and ventral DLPFC and were less apparent in the more posterior and dorsal aspects of DLPFC as well as in PMd. In contrast, action signals are present nearly everywhere perhaps consistent with a global signal^11^, and strongest in PMd. In alignment with the visualization, dPCA loadings for color and mixed selectivity are stronger in anterior DLPFC and in the sulcus (partial correlation, color: *r*_*AP*_ = -0.1168, *r*_*depth*_ = 0.0975, color × action: *r*_*AP*_ = -0.0666, *r*_*depth*_ = 0.08, all p < 0.001). In contrast, dPCA loadings for action are stronger more posterior (partial correlation *r*_*AP*_ = 0.067, p < 0.001) but do not vary as a function of depth (choice: *r*_*depth*_ = -0.016, p > 0.05).

We wanted to further explore this trend as a function of the distinct sub-regions of DLPFC and PMd. We used the spectrolaminar profile^28^ for each session and combined it with recording location to delineate 5 putative sub regions (4 in DLPFC and 1 in PMd, Fig. 5d). For PMd, 8Ad, 9/46A and 9/46d, the averaged spectrolaminar motif demonstrated a “check mark” shape, consistent with recordings from superficial to deep layers, albeit with some differences between PMd and others (e.g., more beta band activity). In contrast, the spectrolaminar motifs of recordings inside the principal sulcus mostly showed an inverted “check mark” shape consistent with recordings starting in deep layers and terminating in superficial layers inside the sulcus (see *Methods: Anatomical localization of recording sites*).

**Figure 5:**
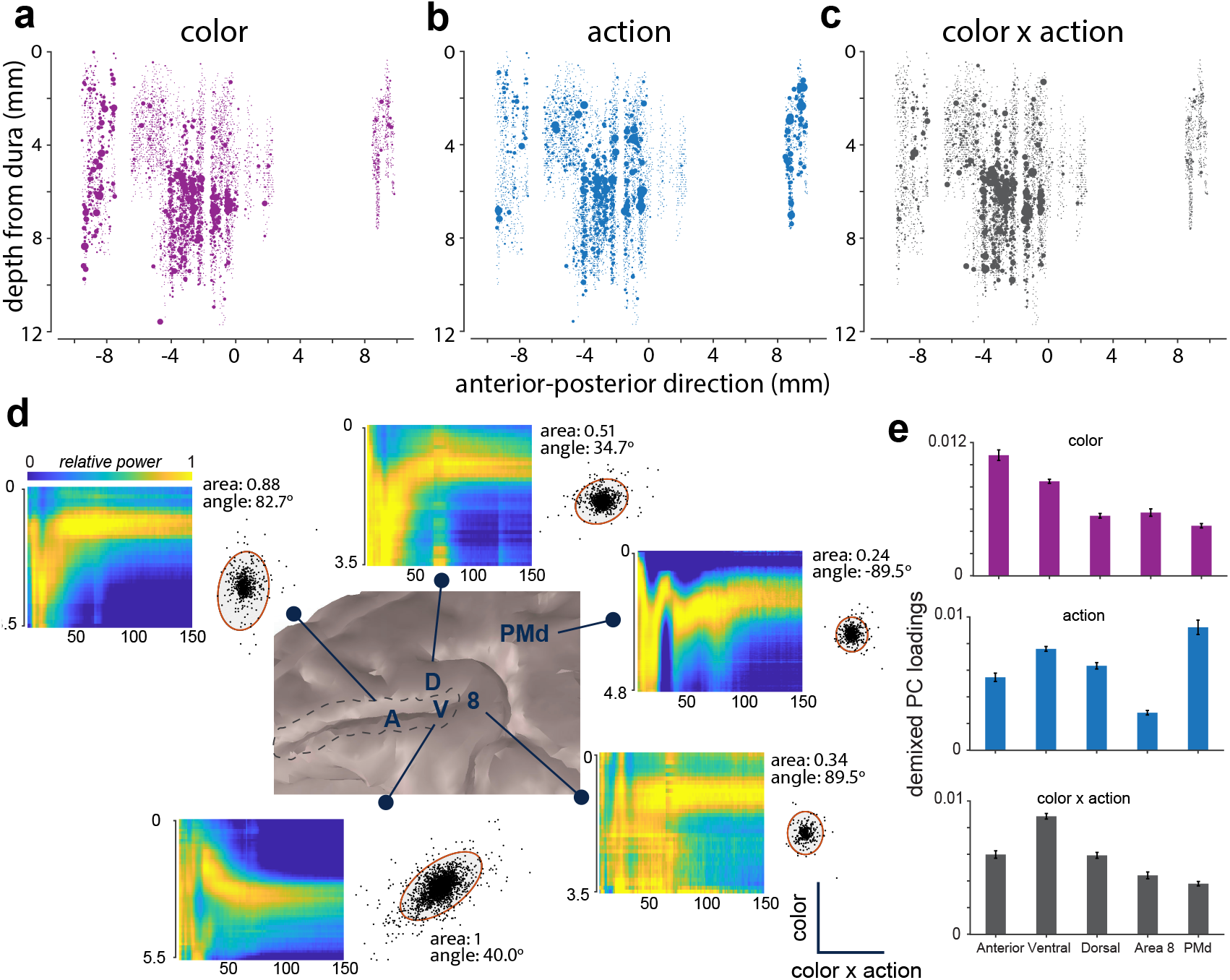
A functional gradient within the frontal lobe. (**a-c**) Average color, action and interaction term (color × action, NMS) dPC loadings for each recorded unit within a time window of [-50, 400] aligned to checkerboard onset. X-axis is the anteriorposterior locations of recordings relative to the start of the principal sulcus with negative numbers indicating anterior and positive numbers indicating posterior. Y-axes denote the depth of the unit. Size of the dot represents the amplitude of dPC loadings; a larger dot represents the unit is more modulated for the corresponding task variable. (**d**) Average spectrolaminar motif of each recorded area and quantification of color and interaction (color × action) selectivity. The scatter plot inset shows the dPC color × action and color loadings for these neurons. We quantified this scatter plot using the area and angle of an ellipse that covered 99% of covariance. The reported elliptical area is the relative area compared to the ellipse for ventral DLPFC. (**e**) Average dPC loadings of each task variable within each area (error bar is SEM). Color signal decreases from anterior to posterior direction. color × action is highest in ventral DLPFC. Action signal is highest in PMd.

For each area, we quantified color and color × action strength using a 99% covariance ellipse estimated from the dPCA loadings (Fig. 5d, scatter plot insets). The ellipse area indicates overall representation strength (Fig. S5g, normalized to ventral DLPFC for comparison), while its angle indicates whether color (90°), interaction (0°), or both (45°) dominate. Fig. S5e quantifies the action and color strength for these 5 brain areas. As illustrated in Fig. 5d, ventral DLPFC demonstrates strong color and color × action selectivity (*angle* = 40°, Fig. S5h). In contrast, anterior DLPFC has the strongest color selectivity (*angle* = 82.7°; bootstrap test between subsample DLPFCv and DLPFCa elliptical angle, p < 0.02). Dorsal and posterior regions have weak color and color × action selectivity (dorsal: *area* = 0.51, bootstrap test between subsample DLPFCv and DLPFCd elliptical area, p < 0.02; area 8: *area* = 0.34, bootstrap test between subsample DLPFCv and DLPFCd elliptical area, p < 0.02). Finally, PMd shows the weakest color and color × action (*area* = 0.24, bootstrap test between subsample DLPFCv and PMd elliptical area, p < 0.02) and strongest action choice selectivity (Fig. S5g, *angle* = 12.7°; bootstrap test between PMd and each of other areas, p < 0.02). Average demixed PC loading (Fig. 5e) and mixed selectivity index (Fig. S5h bottom) for each area also showed a similar pattern as in Fig. 5a-d.

Collectively, these results, enabled by a large multimodal dataset combining electrophysiology, MRI and histological localization, strongly suggest a functional gradient within DLPFC. Mixed selectivity and color representations are concentrated in ventral and anterior DLPFC, decreasing along posterior and dorsal axes toward PMd, while action signals are broadly distributed across frontal cortex and are strongest in PMd.

### Context-specific dynamics or inputs can explain observed neural dynamics

Our results thus far have described that neural responses in DLPFC reflect the XOR computation necessary for flexible decisions and have high-dimensional representations that enable readout of both color and action, and that there is a gradient of color and mixed selectivity. However, two open questions remain. First, what is the underlying mechanism that leads to the XOR computation to be reflected in DLPFC? Second, what are the key factors that generate the observed functional gradients? Our final goal was to use modeling to address these two questions.

A change in context due to a different target configurations could change the dynamics within DLPFC or alter the inputs. Consistent with this view, changes in dynamics, transformations of input, or both can explain neural computations enabling flexible decision making^35;36^. Inspired by these studies, we built variants of a linear firing rate recurrent neural network model^37^ with the same overall dynamical equation for the firing rate **r** (Equation 1, See *Methods: Linear Recurrent Neural Network Model*):

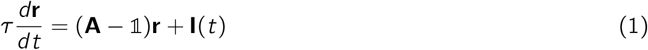

Where **r** is the firing rate of neurons; **A** is the recurrent connectivity matrix; **I**(*t*) is the time varying input; *τ* is the time constant for the dynamics, and 𝟙 is the identity matrix.

We created the following model variants by changing either dynamics or inputs based on context (target configuration in our task): In the switching dynamics (SD) model (Fig. 6a), we implement the nonlinear XOR computation by using different recurrent connectivity matrices (**A**_1_ or **A**_2_, see Fig. S6b) for different contexts, thereby implementing opposite color-to-action mappings. In the flipped input (FI) model (Fig. 6b), context inverts the sign of color inputs while maintaining fixed recurrent dynamics (**A**, see Fig. S6c), reversing the mapping between color and action at the input stage. Finally, in the rotated input (RI) model (Fig. 6c), the dynamics matrix (**A**) is again fixed but context rotates color inputs symmetrically by angle *θ* in opposite directions from color to action axis, directly modulating action through geometric transformation of inputs (see Fig. S6d).

**Figure 6:**
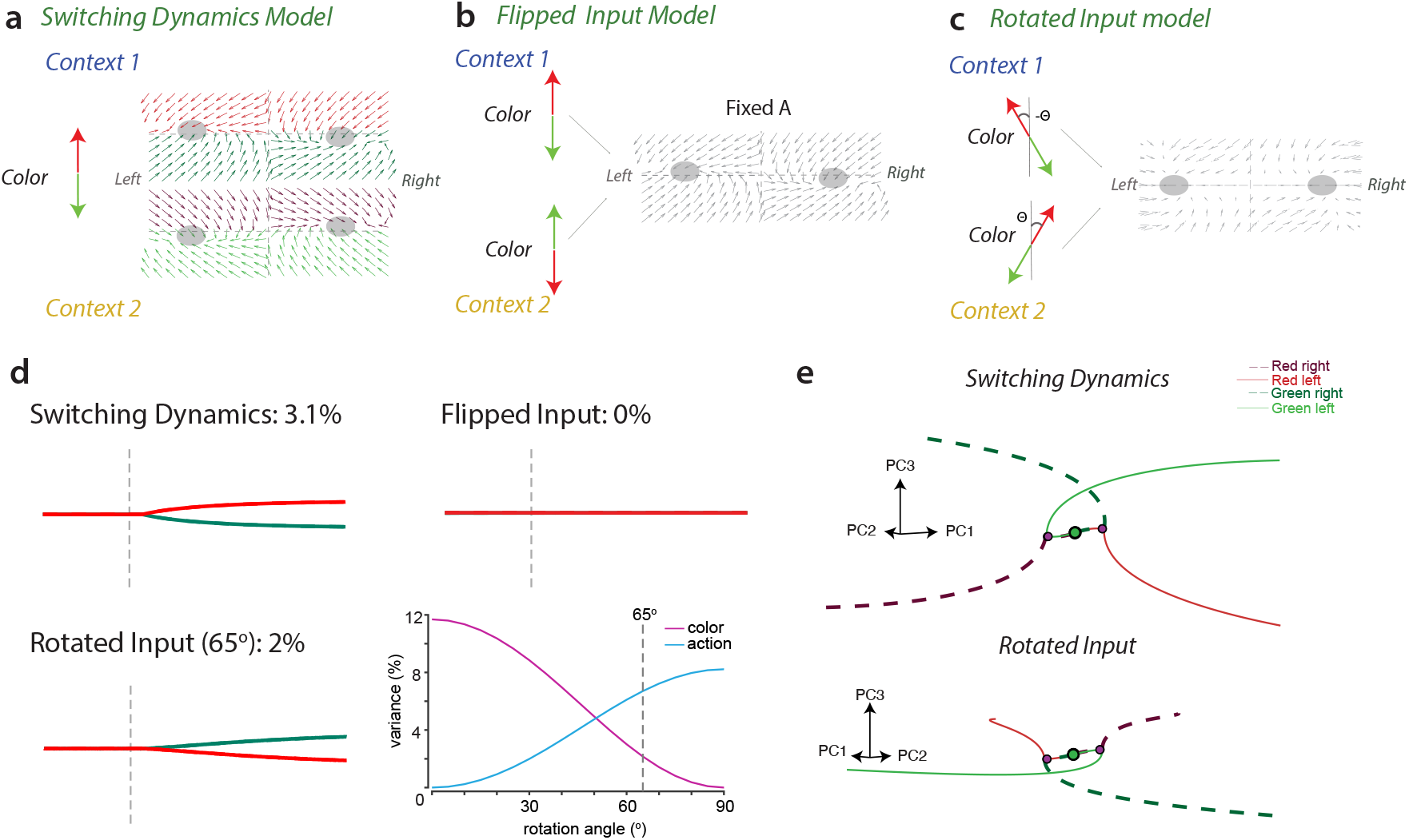
Linear recurrent neural network models with varying dynamics or input transformations can explain patterns in DLPFC. (**a-c**) Diagrams of switching dynamics model (SD), flipped input model (FI) and rotated input model (RI). In the switching dynamics model, the system receives the same color input across contexts but assumes two sets of dynamics (A1 and A2). In the flipped input model, the system maintains same dynamics (fixed A) but contexts flip color inputs to opposite directions. Finally, in the rotated input model, the system again maintains the same dynamics (fixed A); However, the color input is partially aligned with the choice axis and rotates across contexts. (**d**) Projection on to the dPCA color axis for the 3 models. Note, the Switching dynamics, and rotated input models have color variance but the flipped input model does not. Bottom right: we swept rotation angles from 0° to 90° to estimate the best. 65° is chosen to best approximate color and choice variance in DLPFC data. **(e)** PC trajectories projected to PC1, PC2 and PC3 reveal color, action and context signals in both switching dynamics model and rotating input model. Both models capture key features of DLPFC.

We simulated these models and used color and action variance estimated by dPCA to evaluate the model-data match. The SD model predicted reliable color and action variance consistent with dPCA results (Fig. 6d, top left panel). The FI model made a falsifiable prediction: since context perfectly inverts the sign of color inputs, color signals should cancel when combining trials across both contexts, eliminating color variance in dPCA (Fig. 6d, top right panel). This prediction is incompatible with our observed DLPFC data in Fig. 3c, which shows robust color variance. Thus, the FI model cannot fully account for this data. Finally, for the RI model, the rotation angle *θ* affects the variance explained by color and action components in dPCA. At *θ* = 0°, color inputs are identical across contexts, which cancels out action variance across 2 contexts. At *θ* = 90°, inputs project entirely onto the action axis, eliminating color variance while maximizing action variance. Sweeping from 0° to 90° reveals a smooth trade-off: color variance decreases while action variance increases (Fig. 6d, bottom left panel). A rotation of *θ* = 65° replicates the color and action variance observed in DLPFC recordings (Fig. 6d, bottom right panel). Thus, activity of neurons in both the switching dynamics and rotated input model projected on PC1, 2 and 3 (Fig. 6e) are similar to the DLPFC PCA trajectories shown in Fig. 3d. These results again demonstrate that context-dependent sensorimotor mapping in prefrontal cortex can arise through either local dynamic reconfiguration or geometric transformation of sensory inputs^35;36;38^.

Our model in Fig. 6 provides insight into candidate mechanisms in DLPFC. Our second goal was to use modeling to replicate the gradient of selectivity observed in DLFPC and PMd. To this end, we created a variant of our multi-area RNN model from prior work^39^ to model anterior DLPFC (area 1), ventral DLPFC (area 2), dorsal DLPFC/8Ad (area 3) and PMd (area 4). We restricted color and context inputs to areas 1 and 2 and outputs to area 4 (Fig. S7a). We then varied architectural features of the Multi-area RNN model (inputs and feedback connections) and examined which, if any, could replicate the functional gradients observed in the data (Fig. S7b).

Three architectural features were important for replicating the patterns of data observed in Fig. 5. First, feedback connections from areas 3 and 4 to areas 1 and 2 increase the amount of action variance in area 1 and 2 (Fig. S7c). Second, the ratio of color and context inputs to areas 1 and 2 can increase or decrease the gradient. When area 1 receives only color and no context input, area 2 develops stronger nonlinear mixed selectivity and likely becomes the primary area for XOR computation (Fig. S7d), most similar to the gradient observed in Fig. 5. Third, in areas 1 and 2, segregating sensory input units from feedforward units reduces the representations of color and color × action interaction signals in areas 3 and 4 (Fig. S7e-f). Together, the multi-area RNN model provides complementary mechanistic insights into how context modulates sensorimotor transformations in prefrontal cortex and establish testable predictions about the computational roles of feedback connectivity and functional specialization in frontal lobe.

## Discussion

Our goal in this study was to advance our understanding of the neural dynamics that underlie flexible perceptual decisions when the mapping between stimulus and action varies on trial-by-trial basis^3–5;12;20^. We recorded from DLPFC and PMd of monkeys performing a flexible decision-making task that varied the target configuration on a trial-by-trial basis and thus altering the relationship between stimulus and response^3–5^. To solve this task, relevant brain areas need to perform an XOR computation by combining sensory evidence and the target configuration^5^.

Our study provided five key insights. First, we demonstrate distinct neural dynamics and computations in DLPFC and PMd. While both areas exhibit rich decision-related dynamics, firing rates in DLPFC are higher dimensional and robustly encode all task variables, whereas PMd firing rates are lower dimensional and only encodes action choice after it emerges in DLPFC (Fig. 3). Second, DLPFC exhibited a tetrahedron geometry encoding the 4 color/action combinations reflecting a potential solution for the XOR computation necessary for solving this task (Fig. 4). Third, the strength of these representations in DLPFC varies with stimulus difficulty and is weakest for the hardest checkerboard cues. This reduction in mixed selectivity arises because of weaker color representation for more ambiguous checkerboards. In error trials, color signal has a different representation from correct trials and this difference likely leads to delayed action choice signals. Fourth, color and mixed selectivity were strongest in anterior and ventral DLPFC (Fig. 5). Finally, both changes in dynamics and rotation of inputs across target configurations were valid network mechanisms to explain our results (Fig. 6).

These findings provide new insight into the neural dynamics of flexible decision-making and complement prior studies of context-dependent decisions^24;35;40^. A key distinction is that those studies examined tasks where context determines which sensory feature to attend (e.g., color versus motion, or location vs. frequency), whereas our task requires an XOR computation where context reverses the mapping between a sensory feature and motor output—a fundamentally different nonlinear computational problem.

Both PCA trajectories and decoding confirmed that color choice emerged earlier than action choice signals, consistent with recent studies demonstrating that perceptual decisions emerge before motor responses when task design dissociate them from one another^3;4^. Our key contribution is revealing how this dissociation emerges mechanistically: high-dimensional representations in DLPFC unmix color and action before PMd reflects action choice alone. On hard trials and errors, color signals are weaker and action choice signals are correspondingly delayed, demonstrating that the strength of sensory representations directly affects the timing of action selection. Together, these findings demonstrate that perceptual decisions and motor planning involve separable neural representations that can be revealed through appropriate task manipulations.

Our results provide evidence of differences between areas in the frontal lobe of the monkey,^19;33^. Within the areas recorded in this study, nonlinear mixed selectivity and color is strongest in the ventral and anterior DLPFC and decreases posterior towards PMd. In contrast, while action choice is encoded stably across all areas, PMd contains a higher proportion of action-modulated units. Together, these gradients are strong evidence of a shift from abstract high-level computations needed to solve the task to a motion signal that represents the output needed for the animal to perform the task, a pattern that has been suggested by human fMRI studies^18^. These observed gradients align with anatomical studies^41–43^, that suggest systematic differences of inputs to DLPFC. The caudal aspect of DLPFCv receives inputs from the dorsal stream (area LIP) and also from the ventral stream via the VLPFC thus providing inputs that could enable mixed selectivity^44^. In contrast, the rostral aspect of DLPFCv receives stronger color inputs from inferotemporal cortex compared to more posterior sites^34^. These differences in inputs could perhaps explain the results in Fig. 5.

Our findings demonstrate that DLPFC exhibits all the hallmarks of performing the XOR computation necessary to solve this task: it encodes all task variables with nonlinear mixed selectivity, color signals emerge before action signals, and choice representations in DLPFC precede those in PMd. However, these observations alone cannot establish whether DLPFC is the locus where choice signals are computed or whether it reflects computations performed jointly in a distributed network^38;45;46^. As multiple studies and our own modeling show, such task-related dynamics could emerge either from intrinsic recurrent computations within DLPFC or from structured inputs from upstream areas in the dorsal^3;24;47;48^ or ventral stream^49^, and the same is true for the data presented here as well. Resolving this model degeneracy is a critical future direction not only for us but for systems neuroscience. Ultimately simultaneous recordings^50^, pathway specific perturbations, and modeling will help understand how input transformations and local recurrent dynamics combine to orchestrate flexible decisions^51;52^.

## Methods

All data reported in this study were collected under appropriately reviewed and approved protocols. At Boston University experiments for Monkeys T, V, and Z were performed under the approval of the BU Institutional Animal Care and Use Committee, (IACUC, PROTO 201800695). Experiments at Stanford were approved by the Stanford University IACUC (Protocol 8856).

### Subjects

Four adult male macaque monkeys (*Macaca Mulatta*; T, V, Z, and O, Table 1) were trained to perform the behavior task for months, training time varying according to monkey. After the behavior training (as described in^1;2;53^), monkeys underwent sterile surgery where cylindrical head restraint holders (Crist Instrument Co., Inc., Hagerstown, MD, United States) and recording cylinders (round: 19 mm diameter, Crist Instrument Co., Inc. or 33 × 19 mm oval chambers, NaN instruments) were implanted. Round cylinders were placed on surface normal to the cortex and were centered over caudal dorsal premotor cortex (PMdc; +16, 15 stereotaxic coordinates) for monkey T & monkey O. Oval cylinders were placed with similar manner as round cylinders, centered over dorsolateral prefrontal cortex (DLPFC; around +32.5, +15 stereotaxic coordinates) for monkey T, monkey V & monkey Z. The skull within the cylinder was covered with a thin layer of dental acrylic or palacos.

**Table 1:** Subject information.

| Monkey | Sex | Age (years) | Weight (kg) | Chamber Type |
| --- | --- | --- | --- | --- |
| T | M | 15 | 15.0 | Oval/Round |
| O | M | 11 | 15.5 | Round |
| V | M | 14 | 10.9 | Oval |
| Z | M | 14 | 15.0 | Oval |

### Apparatus

Monkeys sat in a customized chair (Synder Chair System, Crist Instrument Co., Inc.) with head fixed. The arm that was not used to respond in the task was gently restrained with a tube and cloth sling. Experiments were controlled and data collected using a custom computer control system (Mathworks’ xPC target/Simulink Realtime and Psychophysics Toolbox, The Mathworks, Inc., Natick, MA, United States). Stimuli were displayed on a monitor approximately 30 cm from the monkey. A photodetector (Thorlabs PD360A, Thorlabs, Inc., Newton, NJ, United States) was used to record the onset of the visual stimulus at a 1 ms resolution. A small reflective spherical bead (11.5 mm, NDI passive spheres, Northern Digital, Inc., Waterloo, ON, Canada) was taped to the middle finger, of the active arm of each monkey; right for T and left for O. The bead was tracked optically in the infrared range (60 Hz, 0.35 mm root mean square accuracy; Polaris system, NDI). Eye position was tracked using an overhead infrared camera with an estimated accuracy of 1° (either ther ISCAN ETL-200 Primate Eye Tracking Laboratory, ISCAN, Inc., Woburn, MA, United States, or our custom built system open source eye tracking system, *openEyeTrack*). To get a stable image for the eye tracking camera, an infrared mirror (Thorlabs, Inc.) transparent to visible light was positioned at a 45° angle (facing upward) immediately in front of the nose. This reflected the image of the eye in the infrared range while allowing visible light to pass through. A visor placed around the chair prevented the monkey from touching the juice reward tube, infrared mirror, or bringing the bead to its mouth.

### Task

Experiments were made up of a sequence of trials that each lasted a few seconds. Successful trials resulted in a juice reward whereas failed trials led to a time-out of 2-4 s. A trial started when a monkey held its hand on a central circular cue (radius = 12 mm). In a subset of sessions, the monkey fixated on a small white cross (diameter = 6 mm) for ∼300-485 ms. Then two isoluminant targets, one red and one green, appeared 100 mm to the left and right of the central hold cue. Targets were randomly placed such that the red target was either on the right or the left trial-to-trial, with the green target opposite the red one. Following an additional center hold period (400-1000 ms) a static checkerboard stimulus (15 × 15 grid of squares; 225 in total, each square: 2.5 mm × 2.5 mm) composed of isoluminant red and green squares appeared superimposed upon the fixation cross. The monkey’s task was to move their hand from the center hold and touch the target that matched the dominant color of the checkerboard stimulus for a minimum of 200 ms (for full trial sequence see Fig. 1). For example, if the checkerboard stimulus has more red squares than green squares, the monkey had to touch the red target in order to have a successful trial. Monkeys were free to respond to the stimulus as quickly or slowly, within an ample ∼2*s* time frame, as they ‘chose’. There was no delayed feedback therefore a juice reward was provided immediately following a successful trial^16^. An error trial or miss led to a timeout until the onset of the next trial.

The checkerboard stimulus was parameterized at 14 levels of red (*R*) and complementing green (*G*) squares ranging from nearly all red (214 *R*, 11 *G*) to all green squares (11 *R*, 214 *G*) (for example stimuli see Fig. 1c). These 14 levels are referred to as signed coherence (*SC*), defined as 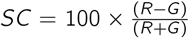 (R: 4%:90%, G: -4%:-90%). Correspondingly there are seven levels of color coherence, agnostic to the dominant color, defined as 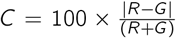 (4-90%). In Fig. 4d-e, we combined easiest 2 color coherence as easy trials; middle 2 coherence as medium and hardest 2 coherence as hard.

The hold duration between the onset of the color targets and onset of the checkerboard stimulus was randomly chosen from a uniform distribution from 400-1000 ms for monkey T, monkey V and monkey Z, and from an exponential distribution for monkey O from 400-900 ms. Monkey O attempted to anticipate the checkerboard stimulus therefore an exponential distribution was chosen to minimize predictability.

### Effects of coherence on accuracy and reaction time (RT)

Behavior was analyzed by estimating psychometric and RT curves for each session and averaging the results across sessions. Behavioral data was analyzed in the same sessions as the electrophysiological data. In total there were 87 sessions for monkey T (136,038 trials) and 66 sessions for monkey O (108,365 trials) for PMd recordings. There were 148 sessions (152,178 trials) for monkey T, 37 sessions (30049 trials) for monkey V and 19 sessions (11421 trials) for monkey Z for DLPFC recordings (we only use monkey Z data in Fig. 5). More details are in Table 2. Both incorrect and correct trials for each *SC* were included for calculating task accuracy and RT. We use psychometric curves to visualize monkeys’ discrimination accuracy. For each given stimulus coherence, we computed the ratio of choosing red on a session-by-session basis and averaged across sessions. Results are displayed in Fig. 1d with error bars denoting SEM and lines between the averages to guide the eyes. Mean RT was calculated per *SC* and averaged across sessions in the same way.

**Table 2:** Sessions, number of trials and units for DLPFC and PMd across all 4 monkeys. Data from monkey Z was only used for Fig. 5.

| Category | Type | DLPFC |  |  |  | PMd |  |  |
| --- | --- | --- | --- | --- | --- | --- | --- | --- |
|  |  | T | V | Z | Total | T | O | Total |
| Sessions | V-probe | 129 | 33 | 19 | 181 | 75 | 66 | 141 |
|  | Neuropixels | 19 | 4 | 0 | 23 | 12 | 0 | 12 |
|  |  | 148 | 37 | 19 | 204 | 87 | 66 | 153 |
| Trials | V-probe | 140,447 | 28,205 | 11,421 | 180,073 | 128,744 | 108,365 | 237,109 |
|  | Neuropixels | 11,731 | 1,844 | 0 | 13,575 | 7,294 | 0 | 7,294 |
|  |  | 152,178 | 30,049 | 11,421 | 193,648 | 136,038 | 108,365 | 244,403 |
| Units | V-probe | 3,577 | 517 | 394* | 4,488 | 546 | 450 | 996 |
|  | Neuropixels | 1,286 | 190 | 0 | 1,476 | 690 | 0 | 690 |
|  |  | 4,863 | 707 | 394 | 5,964 | 1,236 | 450 | 1,686 |

### Electrophysiological recordings

We used single tungsten electrodes, 32-channel V-probes and 45mm Neuropixels 1.0NHP for electrophysiological recordings. Burrholes in the skull were made using handheld drills (DePuy Synthes 2.7 to 3.2 mm diameter). A Narishige drive (MO-972A, Narishige International USA, Inc., Amityville, NY, United States) with a blunt guide tube was placed in contact with the dura. Sharp FHC electrodes (>6M, UEWLGCSEEN1E, FHC, Inc., Bowdoin, ME, United States) penetrated the dura and every effort was made to isolate, track, and stably record from single neurons. 180 *µm* thick 32-electrode linear multi-contact electrode (U-probe, Plexon, Inc., Dallas, TX, United States); interelectrode spacing: 100, contact impedance: 100k) and 384-channel, 110 *µm* thick, 45mm Neuropixel NHP 1.0 probes (IMEC, Inc., Leuven, Belgium) were performed similarly to single electrode recordings with some modifications. We mounted the probes on a A NaN CMS drive (NAN Instruments, Israel) with a sharpened guide tube aided in slow penetration (3-5 *µm*/s). V-probe and Neuropixel penetration was stopped once a reasonable sample of neurons was acquired. Neural responses were allowed to stabilize for 45-60 minutes before normal experiment began. Additionally, lowering the electrode necessitated careful observation to ensure the electrode did not bend or break at the tip, or excessively dimple the dura.

### Unit classification

For 32-channel V-probe, single neurons were delineated online by the ‘hoops’ tool of the Cerebus system software client (Blackrock Microsystems, Salt Lake City, UT, United States). As for V-probe channels capturing multiple neurons, different gates were applied manually if the waveforms were separable during recordings.

For Neuropixel recordings, we used Kilosort4 for post-hoc spike sorting. Then the results were passed to phy for manual curation. Neural signals were selected from noise based on waveform with waveform amplitude greater than 10 and spatial decay greater than 0.01 and less than 0.7. We selected single units from multi-units based on auto-correlogram and waveform shapes. Units are merged if they have similar cross-correlogram, PC projections and cosine template similarity greater than 0.9. Units are split if they have more than one clusters in PC projections. In Fig. 1h (right), we plotted the relative recording depth of each averaged unit waveform. The waveform is classified as positive (blue color) if absolute value of peak is greater than absolute value of trough Otherwise, it’s classified as negative waveform (red color).

### Peri-stimulus time histogram (PSTH)

We calculated peri-stimulus time histograms with the following procedure: 1) We first binned spike times for each trial at 1 ms resolution in a time window from -200ms to 600ms aligned to targets (targets epoch) and from -800ms 800ms aligned to checkerboard onset (checkerboard epoch), respectively. 2) We then convolved the spike train with a Gaussian kernel (*σ*=25ms) to estimate the instantaneous firing rate for a trial. 3) Finally, in target epoch, firing rates were averaged across trials according to each of the 2 target configurations, resulting in 2 trial-averaged PSTHs. In the checkerboard epoch, firing rates were trial-averaged according to chosen color (red or green) and action choice (left or right) in checkerboard epoch. As a result, we derived 4 trial-averaged PSTHs: red left (RL), red right (RR), green left (GL), green right (GR).

### Selection of modulated units

We selected the modulated units based on task variable-averaged PSTH. Chosen units were included if their firing rates are higher than 5 spikes/s, and have modulated activity to any task variable (including increase or decrease in firing rates) in at least one task epoch. We included multi-units as well since they gave us additional power for our analyses and prior work has shown that the inclusion of multi-units does not distort recovery of low-dimensional dynamics from neural activity^54^. In total, for DLPFC, we selected 4863 units from monkey T, 707 units from monkey V and 394 units from monkey Z. for PMd, we selected 1236 units for monkey T, 450 units from monkey O. Table 2 provides a summary of modulated units included in our analysis.

### Population dynamics

#### Principal Components Analysis (PCA)

##### Condition-averaged firing rates

For every unit **n** in total units number **N**, we averaged the single-trial firing rate by chosen color **S** (green or red) and action choice **D** (left or right). As a result, a 4D firing-rate tensor **X**^N×S×D×T^ was created as input to demixed principal component analysis algorithm. For PCA, firing matrix was calculated separately in target period. We averaged firing rates based on two target configurations: **X**^1×C×T^. To ensure the dimensions in target segment same as that in checkerboard segment, the missing dimension is filled with NaN.

For neural data, due to the stochasticity in task design, there is a trial-by-trial difference in interval between target and checkerboard onset (TC interval). The reaction time (from the checkerboard onset to monkey’s hand movement initiation) also varies for each trial. To align time events across trials, we chose a data segment [−100*ms*, 700*ms*] aligning to target and a data segment [0*ms*, 500*ms*] aligning to checkerboard and concatenated them together.

##### Normalization

Due to the heterogeneity in firing rates, PCA results might be dominated by units with high firing rates. To equalize the contribution of each unit in PCA, for each unit **n** ∈ N, 99 percentile of max firing rate **x**_n,99_ was calculated over all color, action conditions and time point. Then we normalized the data by dividing firing rate by 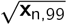. The condition independent signal is another source that explains substantial amount of population variance other than task-related signal. Before conducting principal component analysis (PCA), we calculated the average firing rate of each single unit **X**^1×1×1×T^ over both color (red vs green) and actions (left vs right) conditions and subtracted this condition independent signal from the original data **X**^1×S×D×T^

PCA was used to examine firing rate variance in the recorded neural population. PCA reveals dimensions that explain a large percentage of the data while making few assumptions about the underlying structure of the data. The firing rate matrix **X**^N×S×D×T^ was reorganized into 2D matrices **X**^N×SDT^ appropriate for PCA. Then, PCA projects high-dimension data into low-dimensional axis which maximize the variance in the data by minimizing the loss function:

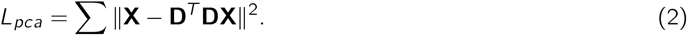

**X**^N×T^ is high-dimension raw data and **D**^N×N^ is the decoding matrix. The low-dimension trajectories **X**^M×T^ (*M* < *N*) calculated by multiplying first *M* rows of **D** by **X**.

For PMd data, principal axis 1, 2 and 3 were plotted which maximize the variance explained. For DLPFC data, principal axis 1, 3 and 4 were plotted to provide a better visualization of the evolution of neural trajectories.

#### Test dimensionality

To test whether DLPFC neural activity has higher dimension than PMd, we subsampled 50 to 1600 units with step size of 150 units (*n* ∈ {50, 200, 350, …, 1600}) randomly from raw data and condition independent signal removed data and performed PCA. We repeated this process for 100 times and reported numbers of PCs required to explain 90% variance. We plotted the mean PC number; error bar is standard deviation of the mean obtained through bootstrapping. We calculated the difference in the dimensionality between DLPFC and PMd and counted the number of instances where the dimensionality of PMd exceeded that of DLPFC. If dimensionality of DLPFC never exceeded that of PMd, then we report the minimum p-value for 100 bootstraps (2/100).

#### Demixed principal components analysis

We performed demixed principal components analysis (dPCA) to quantify variance of neural activity explained by task variables. dPCA is a dimensionality reduction technique that projects data onto task variable related axis while preserving overall variance by minimizing a loss function:^32^.

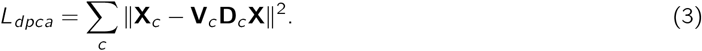

Here, **X**_*c*_ refers to condition-averaged firing rates, having the same shape as **X** ∈ ℝ^*N*×*cT*^, but with the entries replaced with the condition-averaged response. The aim is to recover (per dPCA condition *c*) a **V**_*c*_ and **D**_*c*_ matrix. **V**_*c*_ is constrained to have orthonormal columns, while **D**_*c*_ is unconstrained. The number of columns of **V**_*c*_ and rows of **D**_*c*_ reflects the number of components one seeks to find per condition. The column of **V**_*c*_ is the loading of each neuron to the targeted dPC axis, which reflects how much the demixed axis contributes to each neuron. In dPCA performed in Fig. 3c, we used all correct trials so chosen color is equivalent to stimulus color. For other dPCA analysis, we used all trials and chosen color.

We performed a permutation test to investigate whether DLPFC color and target configuration variance is significantly above zero. For each session, we shuffled the color label and created the 4D firing-rate tensor and concatenated each session. We repeated this process for 100 times to create a null distribution. We defined p-value as the probability of color variance of this null distribution exceeded observed DLPFC color variance. We repeated the same procedure for target configuration.

To test whether color and target configuration variance explained in DLPFC is higher than PMd, we compared the dpc variance difference between subsampled DLPFC and PMd datasets and a surrogate dataset. We created a surrogate dataset by sub-selecting 800 neurons from DLPFC **X**_800pfc_ and PMd **X**_800pmd_ dataset each, combined them and randomly split in half as group A **X**_A_ and B **X**_B_. We calculated the dpc variance explained difference between **X**_A_ and **X**_B_ (*d*_*null*_), and between **X**_800pfc_ and **X**_800pmd_ (*d*_*sample*_) with respect to each task variable. We repeated this process for 1000 times. To test whether color and target configuration variance in DLPFC is higher than PMd, p-value is defined as the proportion of *d*_*sample*_ −*d*_*null*_ below 0. To test whether action variance in DLPFC is lower than PMd, p-value is defined as the proportion of *d*_*sample*_− *d*_*null*_ above 0. The minimal p value we can get is 0.002 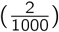.

#### single-unit encoding of task variables

We quantified single unit modulation of task variables with two approaches.

#### 2-way ANOVA

First, we used a 2-way ANOVA to identify the percentage of modulated neurons. For each neuron, we calculated binned spike counts using a 50 ms sliding window with 5 ms steps for each trial *m* ∈ *M* to generate a spike count matrix **X** ∈ ℝ^*M*×*T*^, where *M* is the number of trials and *T* is the number of time bins. We then performed a 2-way ANOVA with main effects of color choice and action choice, and an interaction term (color × action) at each time window. We calculated and plotted the percentage of neurons with significant main effects and interaction terms at each time window using *p* < 0.01. Mathematically, before checkerboard onset, the color × action interaction is equivalent to target configuration, as color and action are perfectly correlated within each configuration. After checkerboard onset, this interaction reflects nonlinear mixed selectivity for color choice and action choice.

#### Modulation Index

Second, to quantify the strength of the encoding of these task variables, we computed modulation indices for each unit at each time window.

For instance, the modulation index for action is defined as:

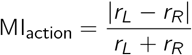

*r*_*L*_ and *r*_*R*_ are trial-averaged spike counts for left and right choices, respectively. Similarly, for color, we have

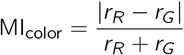

For the interaction term, we calculated the index by using the red&left and green&right trials.

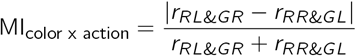

We plotted the average modulation index at each time bin across all units in DLPFC and PMd, respectively.

#### Joint PCA of DLPFC and PMd

We analyzed the neural activity in a window from -100ms to 300ms relative to checkerboard onset. For each unit, we calculated the trial-averaged binned firing counts *X*^1×*S*×*D*×*T*^ of each color *S* and action *D* from the spike counts matrix *X*^*M*×*T*^ (M trials by T time points, mentioned in previous method section). We concatenated these condition-averaged responses across time into a single vector of length *SDT* = *S*×*D*×*T*. This data is normalized in the same manner as PCA but the condition averaged signal is not removed. We combined the binned firing counts from all PFC units 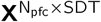 (*N*_*pf c*_ = 5, 570) and PMd units 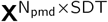 (*N*_*pmd*_ = 1, 686) into a single data matrix 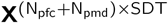 (n = 7256).

We applied PCA to the concatenated population activity to identify low-dimensional subspaces that captured shared variance across both regions. The PCA decomposition yielded a coefficient matrix D (7,256 units × k components). Each element *d*_*ij*_ represents the loading of unit *i* ∈ 7256 onto the principal component *j* ∈ *k*. To examine each region’s contribution to this joint latent space, we extracted region-specific coefficient matrices by partitioning D according to the regional identity of each unit: *D*_*pf c*_ corresponded to rows 1–5,570 (PFC units) and *D*_*pmd*_ corresponded to rows 5,571–7,256 (PMd units). We then projected each region’s activity separately using these region-specific loadings:

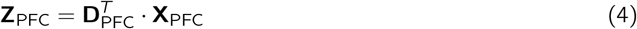

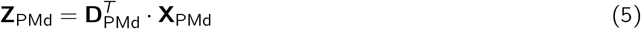

where **D**_PFC_ ∈ ℝ^5570×*k*^ and **D**_PMd_ ∈ ℝ^1686×*k*^ are the loading coefficients for PFC and PMd units respectively from the joint PCA, and **X**_PFC_ ∈ ℝ^5570×*SDT*^ and **X**_PMd_ ∈ ℝ^1686×*SDT*^ denote the normalized firing rate matrices for each region. After this step, we reshape **Z**_PFC_ and **Z**_PMd_ to dimension: **k** × **S** × **D** × **T**. In Fig. 4a and Fig. S3a, **S** is red vs green and **D** is left vs right. We visualized these low dimensional trajectories (projected to PC3 and PC7 in Fig. 4a; projected to PC4 and PC5 in Fig. S3a) of each color/action combinations: red left (RL), red right (RR), green left (GL) and green right (GR) at each time point.

#### Quantify task variable separation in PC space

To quantify these separations in PC space, we choose top 10 PC (*>* 90% variance explained) and computed euclidean distance between each task variables. At a specific time point, we have low-dimensional projection of DLPFC activities: **Z**_RL_, **Z**_RR_, **Z**_GL_, **Z**_GR_ as a 10 dimensional vector for each conditions. Average color separation is defined as:

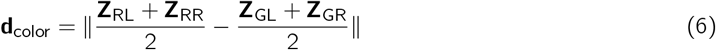

Average color x action interaction term separation is defined as:

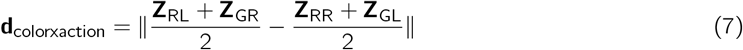

Average action separation is defined as:

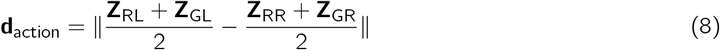

Average nonlinear choice separation is defined as:

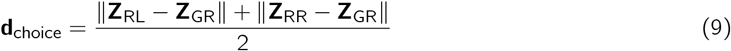

We performed these calculations at each time point to plot the separation of task variables as function of time. Same calculation was applied for PMd low-dimensional data.

##### Bootstrap test for significance

To test the significance of differences in task variable separation (Fig. 4b), we used a bootstrap procedure. We randomly subsampled DLPFC neurons to match the PMd dataset size. For each bootstrap iteration, we performed PCA jointly on the subsampled DLPFC data and complete PMd dataset. For each task variable, we computed the separation difference between DLPFC and PMd (*d*_DLPFC_ − *d*_PMd_) and averaged this difference within a 150-250 ms window after checkerboard onset. We repeated this procedure 100 times and calculated the p-value as the proportion of bootstrap samples where DLPFC separation was less than or equal to PMd separation. When we found only zero bootstrap samples meeting this criterion, we report the minimum two-tailed p-value of *p* < 0.02 (≈ 2/101). The same procedure was used to compare DLPFC nonlinear choice separation versus pure choice separation.

##### Volume and Surface area

To quantify the nonlinear mixed selectivity, we calculated tetrahedron volume and surface area embedded in 10 dimensional PC space bounded by 4 color/choice combinations. The surface area S was computed as the sum of the areas of the four triangular faces:

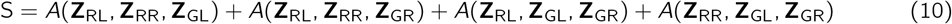

For each triangular face with vertices **a, b, c**, the area was calculated using the Gram Matrix method:

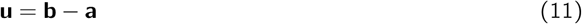

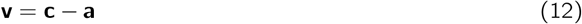

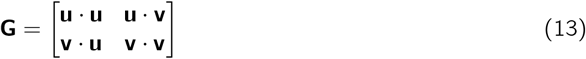

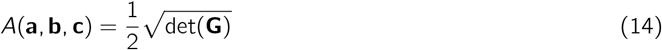

The volume of the tetrahedron was computed using the Cayley-Menger determinant via the Gram Matrix. We constructed three edge vectors from vertex **Z**_RL_:

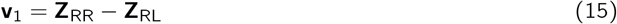

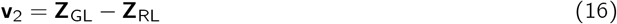

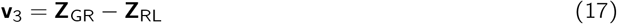

The Gram Matrix captures all pairwise inner products between these vectors:

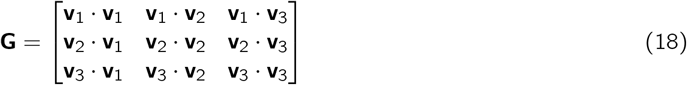

The volume of the tetrahedron is:

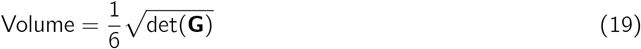

##### Bootstrap test

To test the significance of differences in volume and surface area between DLPFC and PMd, we used the same bootstrap procedure as for task variable separation (Fig. 4b): subsampling the DLPFC dataset to match PMd size, computing volume or surface area for both subsampled DLPFC and PMd, and calculating the proportion of bootstrap iterations where DLPFC was less than or equal to PMd. We used 100 bootstrap iterations. When zero iterations showed DLPFC ≤ PMd, we report the minimum two-tailed p-value of *p* < 0.02 (2/100 bootstrap iterations).

In Fig. 4d and Fig. S3b-c, we performed and plotted task difficulty dependent PCA at multiple time points. We manually separated our 7 task difficulties into easy (2 easiest color coherence, red: 214, 190; green: 11,45), medium (2 middle color coherence, red: 158, 147; green: 67,58) and hard (3 hard color coherence, red: 135, 124, 117; green: 90, 101, 108). We averaged the firing rates for these task difficulties and created a firing rate tensor **X**^*N*×*S*×*D*×*T*^ where *S* = 6 (3 red difficulty and 3 green difficulty). Then we reshape the tensor to **X**^*N*×*SDT*^ and performed PCA. For better visualization, we subtracted the condition average signal after PCA.

#### Single-trial decoding of color, mixed selectivity, and action choice

We also wanted to complement insights from dimensionality reduction on trial-averaged firing rates and assess and understand computations in DLPFC and PMd at the single-trial level. For this goal, we used decoding to estimate the time evolution of various task variables.

We only used Neuropixels recordings from DLPFC (n=23, 19 from T and 4 from V) and PMd (n=12 from T). For Neuropixels recording sessions (n = 23), we built a decoder to predict task variables with firing rates. For each session, we built the spike counts tensor *X* ∈ ℝ^*M*×*N*×*T*^, where *M* is the number of trials, *N* is the number of units, and *T* is the number of 50ms time bins. For each time point *t* ∈ *T*, we trained a L2-regularized logistic regression classifier with 5-fold cross validation to predict the average accuracy of the target binary classes.

In Fig. 4c, we reported the nonlinear choice and pure choice decoding accuracy as a function of time. Nonlinear choice, also known as context-dependent choice, is the separation between left and right with respect to each target configuration. In target configuration 1, context-dependent choice separates red left (RL) and green right (GR). In target configuration 2, context-dependent choice separates red right (RR) and green left (GL). We built two decoders *d*_*RL*|*GR*_ and *d*_*RR*|*GL*_ using equalized trials. We reported the average accuracy of these 2 decoders as nonlinear choice accuracy. For pure choice, we chose the same trials as each of the contextdependent choice decoder *d*_*L*|*R*_ to predict left vs right choices and reported the accuracy as pure choice accuracy. To test whether the decoding accuracy between nonlinear choice and pure choice are significantly different at each time bin, we performed a paired t-test for each bin.

##### Latency

To estimate the latency when decoding accuracy for pure choice and nonlinear choice are significantly above chance, we performed a one-sample t-test comparing accuracy at each time point after checkerboard onset to the average accuracy during the baseline period (before checkerboard onset). We identified sequences of consecutive time bins that were significantly above baseline (*p* < 0.01) for at least 10 consecutive bins (50 ms with 5 ms bin steps) and report the first time bin in such a sequence as the latency. Bootstrap confidence intervals for latency estimates were computed by resampling sessions with replacement (5000 iterations) and repeating the latency estimation procedure on each bootstrap sample.

In Fig. 4e, we built 3 nonlinear choice decoders *d*_*easy*_, *d*_*medium*_, and *d*_*hard*_ using easy, medium and hard trials, respectively. We plotted the time-dependent nonlinear choice decoding accuracy for easy, medium and hard difficulties. To test whether differences of nonlinear choice decoding accuracies are significant, we performed an one-way ANOVA on each task difficulty’s averaged decoding accuracy across a [100*ms*, 300*ms*] window after checkerboard onset. In Fig. 4f-g, we also trained 3 decoders (with only correct trials) to predict color and choice using easy (2 easiest color coherence, red: 214, 190; green: 11,45), medium (2 middle color coherence, red: 158, 147; green: 67,58) and hard (3 hard color coherence, red:135, 124, 117; green: 90, 101, 108) trials. In this analysis, we chose sessions (n = 16) with at least 5 error trials for each of the color/action combination (at least 40 error trials in total). To assess whether correct and error trials share the same subspace, we trained a decoder using hard correct trials, then tested the decoding accuracy with equal number of hard correct and wrong trials.

To test whether the color decoding accuracy changes with task difficulty, we performed an one-way ANOVA on the averaged color decoding accuracies across a [100*ms*, 300*ms*] window after checkerboard onset. To test if color was stronger than action choice, we chose the average decoding accuracy in [100*ms*, 200*ms*] window after checkerboard onset and performed a paired t-test (n = 23). To test the decoding accuracy difference between correct and error trials, we chose the averaged accuracy across a time window of [200*ms*, 500*ms*] after checkerboard and performed a t-test between hardest correct trials and error trials accuracies (n =16). Finally, to estimate the latency for color and action decoding, we used the same approach as for the nonlinear and pure choice.

Finally, in Fig. S4b, we built 6 decoders to separate each pairs of color/action combination. They are *d*_*RL*|*RR*_, *d*_*RL*|*GL*_, *d*_*RL*|*GR*_, *d*_*RR*|*GL*_, *d*_*RR*|*GR*_, *d*_*GL*|*GR*_. We plotted the decoding accuracy as a function of time and calculated *R*^2^ of each pairs of these decoding accuracies.

#### Anatomical localization of recording sites

We estimated each burrhole’s center location based on structure MRI of monkey T. During MRI scan we put Vitamin E filled capillary tube in a grid hole location representing the center of burrhole 3. Vitamin E filled capillary tube left a trajectory in MRI scan. We segmented brain and visualized the location of burrhole 3 with software 3DSlicer (See Fig. S1a-b). Based on this method, posterior edge of burrhole 3 is identified as the posterior end of principal sulcus, served as zero point in anterior-posterior (AP) direction. We calculated the AP coordinate of each burr hole center relative to this landmark. We documented each session’s burr hole identity after experiments. For monkey V and Z, we validated AP locations post-mortem by extracting the brain and measuring the AP distance from each burrhole center to posterior end of principal sulcus (Fig. S1c-d). Then we estimated each session’s recording AP location as burr hole center with a uniformly distributed jitter of ±1 mm. We calculated depth of each unit from dura by combining each day’s insertion depth with probe channel spacing (V-probe: 100 *µ*m; Neuropixel: 20 *µ*m).

To assess each recording unit’s modulation to color, action and their interaction, we combined all DLPFC and PMd binned firing counts data (see *Methods: Population dynamics*) from -100ms to 400ms around checkerboard onset and performed dPCA. We extracted each unit’s loading on first dPC color, action and color × action axis. In Fig. 5a-c, marker size represents loading magnitude (each dPC loading is scaled by 400 for visualization). To examine whether task modulation is correlated to anatomical position, we performed partial correlation analyses relating dPC loadings to recording depth and A/P location. For color, color x action, and action, we computed two partial correlations: (1) the correlation between dPC loadings and depth while controlling for A/P, and (2) the correlation between dPC loadings and A/P while controlling for depth. In Fig. S5a-c, we also reported the dpca loadings without normalization (each dPC loading is scaled by 400). In Fig. S5d-f, we reported averaged effect size (over a time window of [100, 400] aligs to checkerboard onset) quantified as Cohen’s d as each unit’s mean firing rate differences between conditions. (each Cohen’s d is scaled up by 70 for visualization).

In Fig. 5d, we classified each recorded unit to one of the 5 areas (anterior, dorsal, ventral, area 8 and PMd). First, based on recording AP locations, we classified all Monkey V units as anterior, all monkey T and Z posterior recordings as area 8. We referred to LFP spectrolaminar analysis to differentiate monkey T and Z superficial vs ventral recordings. We inferred laminar locations from spectral power gradients in beta and gamma band. Recordings traversing superficial-to-deep layers (e.g., superficial recordings area 8) exhibited a characteristic increase in gamma power (75–150 Hz) and decrease in beta power (10–19 Hz) with depth, thus forming a “check mark” shape. Recordings traversing deep-to-superficial layers (e.g., penetrating from white matter to 9/46d) showed inverted spectral gradients (inverted “check mark” shape). For penetrations spanning multiple areas, we identified transitions between spectrolaminar motifs and segmented areas manually.

After we segmented recording areas, we plotted average spectrolaminar profile of each area. We estimated putative layer 4 channel using FLIP algorithm, the channel where gamma and beta band power are equal. For recordings with LFP 60Hz noise contamination or cover multiple cortical areas, layer 4 channel was estimated based on shape of spectrolaminar profile. We aligned and concatenated all single-day spectrolaminar profile to putative layer 4 channel and calculated the mean profile with nanmean function in matlab. For each segmented recording area, we generated the 2D scatter plot of each unit’s dPC loadings for color x action versus color. We plotted a covariance ellipse that covers 99 percentage of the covariance (99 percentile confidence interval). Ellipse angle is defined as the angle between major axis and minor axis. Relative area is the proportion of the ellipse area in each area compared to ventral DLPFC. In Fig. S5g, we plotted the 2D scatter plot of color and action dPC loadings. For each task variable, we reported the average dPC loadings of all units in each area (error bar: SEM) as a bar graph.

We defined a mixed selectivity index 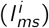 to quantify each unit’s mixed selectivity of multiple task variables:

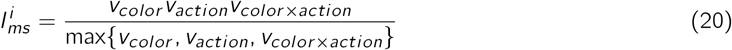

where *v*_*color*_, *v*_*action*_, *v*_*color*×*action*_ are dPC loadings of each unit on top color, action and color x action axis. we performed bootstrap test to examine whether that observed covariance ellipsis angle and area difference shown in Fig. 5d are artifacts of substantially more units in DLPFCv. We randomly sampled 1000 units from DLPFCv *X*_1000*pf cV*_, merged these with all data from other areas, computing each unit’s color, action and color x action dPC loadings, and calculated the color&action and color&color x action covariance ellipse angle and absolute area. We repeated this process for 100 times and generated a distribution of these 2 metrics. The mean areas are: DLPFCa (0.002 ± 0.00011), DLPFCv (0.0032 ± 0.00011), DLPFCd (0.0017 ± 0.000068), area 8 (0.0011 ± 0.000043), and PMd (0.0008 ± 0.00002). Mean angles were: DLPFCa (80.45° ± 0.46°), DLPFCv (35.40° ± 1.29°), DLPFCd (24.39° ± 1.40°), area 8 (88.85° ± 0.64°), and PMd (88.76° ± 1.06°). The small standard deviations confirmed the observed elliptical area and angle differences are not due to sample size imbalance. We test the angles difference between DLPFCv and DLPFCa; area difference between DLPFCv and DLPFCd, DLPFCv and area 8, and DLPFCv and PMd. For color&color x action covariance, we test the ellipsis angle difference between PMd and each of other areas.

#### Linear Recurrent Neural Network Model

Our neural data suggest that neural dynamics in DLPFC leads to neural representations that covary with target configuration, color choice, and action choice. To understand the underlying mechanism, we constructed linear recurrent neural network models consisting of *N* = 100 neurons with the following dynamical equation:

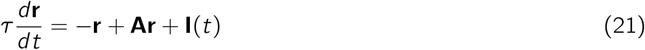

where **r** ∈ ℝ^*N*^ represents the firing rates of the neurons; *τ* = 50 ms is the time constant; **A** ∈ ℝ^*N*×*N*^ is the connectivity matrix; and **I**(*t*) is the input. All simulations used a time step of *dt* = 1 ms over a total trial duration of *T*_total_ = 700 ms.

##### Connectivity matrix

Inspired by Murray et al.^37^, we constructed a connectivity matrix **A** with two components:

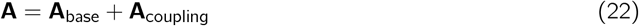

Following Murray et al.^37^, the general form of the connectivity matrix is:

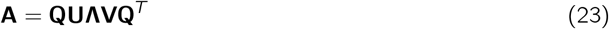

where **Q** is a rotation matrix, **Λ** is a diagonal matrix of eigenvalues, and **U** and **V** are related by **V** = **U**^−1^. This general form allows for non-normal dynamics where left and right eigenvectors differ. For simplicity, we set **U** = **V** = **I**, yielding the symmetric form:

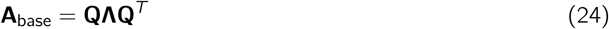

We generated the orthonormal basis **Q** ∈ ℝ^*N*×*N*^ via QR decomposition of a random Gaussian matrix (random seed = 42). This basis defined four principal computational dimensions: context (dimension 1), color (dimension 2), choice (dimension 3), and condition-independent activity (dimension 4) (Fig. S6a left). The diagonal matrix **Λ** determined the temporal stability of each dimension, with eigenvalues set to: context (Λ_11_ = 0.2), color (Λ_22_ = 0.15), choice (Λ_33_ = 0.9), and condition-independent signal (Λ_44_ = 0.75). Remaining eigenvalues were set as background signals:

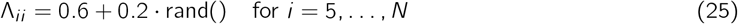

This eigenvalue structure ensures that context information is weakly maintained, color signals are transient, action representations are stable, and condition-independent activity provides persistent background activity on top of which task-related signals unfolded.

In Fig. S6a right, the second component is **A**_coupling_, a rank-1 connectivity matrix that couples color and action dimensions. The coupling matrix was defined as:

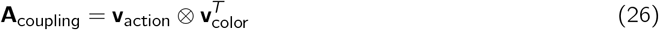

where **v**_action_ = **Q**(:, 3) and **v**_color_ = **Q**(:, 2) are the action and color loading vectors from **Q**, and ⊗ denotes the outer product. This rank-1 coupling creates a directed mapping from the color dimension to the action dimension, enabling color information to drive action selection.

##### Input structure

Following Murray et al.^37^, for the network to perform the XOR computation, inputs must project onto their corresponding dimensions in the connectivity matrix. Specifically, color inputs must align with the color dimension (column 2 of **Q**) and context inputs must align with the context dimension (column 1 of **Q**). Importantly, each neuron receives a heterogeneously weighted version of these inputs, determined by randomly drawn tuning weights ***η***, ensuring diversity in single-neuron selectivity profiles. We implemented this as follows.

The total input to the network at time *t* has three components:

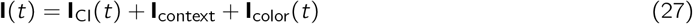

The context input **I**_context_ is present throughout the trial:

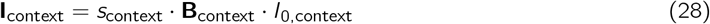

where *s*_context_ ∈ {−1, +1} indicates target configuration (T_1_: +1, T_2_: -1). **B**_context_ is the context input weight vector. **B**_context_ = **Q**(:, 1) ⊙ ***η***_context_ projects onto the context dimension with neuron-specific weights ***η***_context_ ∼ 𝒩 (0, 1) (normalized to unit mean), and *I*_0,context_ = 0.7.

The color input **I**_color_(*t*) is presented at *T*_color_ = 230 ms:

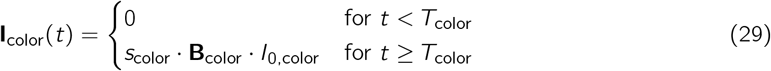

where *s*_color_ ∈ {−1, +1} represents color (green: -1, red: +1). **B**_color_ is the color input weight vector. **B**_color_ = **Q**(:, 2) ⊙ ***η***_color_ projects onto the color dimension with neuron-specific weights ***η***_color_ ∼ 𝒩 (0, 1), and *I*_0,color_ = 0.6.

A ramping condition-independent input **I**_CI_(*t*) drives background dynamics:

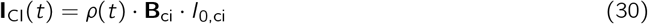

where *ρ*(*t*) = 0.8 *·* (*t/T*_total_), **B**_ci_ = **Q**(:, 4) ⊙ ***η***_ci_ with neuron-specific weights ***η***_ci_ ∼ 𝒩 (0, 1), and *I*_0,ci_ = 7.0. We simulated four experimental conditions corresponding to the factorial combination of context (T_1_ vs. T_2_) and color (red vs. green):

1. context: T_1_ (*s*_context_ = +1), color: red (*s*_color_ = +1), action = Left (*s*_choice_ = −1)
2. context: T_1_ (*s*_context_ = +1), color: green (*s*_color_ = −1), action = Right (*s*_choice_ = +1)
3. context: T_2_ (*s*_context_ = −1), color: red (*s*_color_ = +1), action = Right (*s*_choice_ = +1)
4. context: T_2_ (*s*_context_ = −1), color: green (*s*_color_ = −1), action = Left (*s*_choice_ = −1)

Each condition was simulated independently with the same network parameters and initial conditions (**x**(0) ∼ 0.01 *·* 𝒩 (0, **I**)).

We implemented three models that differ in how context (target configuration) modulates the color-to-action transformation:

##### Switching Model

As illustrated in Fig. 6a and Fig. S6b, in the switching model, the sign of **A**_coupling_ reverses based on the context. The full connectivity matrices are:

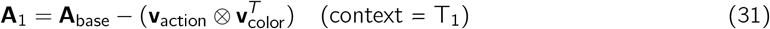

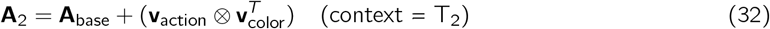

##### Flip Input Model

In the flip input model, the connectivity matrix remains fixed as equation (22), but the color input is flipped in T_1_ and T_2_ trials, respectively (Fig. 6b and Fig. S6c):

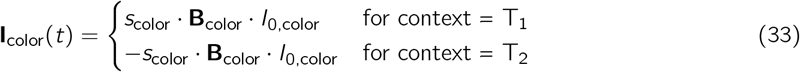

where *s*_color_ ∈ {−1, +1} indicates red or green color, and **B**_color_ ∈ ℝ^*N*^ is the color input weight vector. This implements an additive input transformation where context determines whether the color signal is inverted before entering the network.

##### Rotated Input Model

In the rotated input model (Fig. 6c and Fig. S6d), the connectivity matrix remains fixed as in equation (22), but the color input is geometrically transformed by rotating it toward the action dimension. Critically, this requires an input direction **B**_action_ = **Q**(:, 3) ⊙ ***η***_action_ that is aligned with the action dimension in the connectivity matrix. The rotated input is:

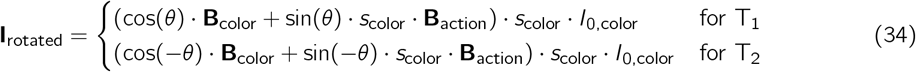

where *θ* is the rotation angle and *s*_color_ ∈ {−1, +1} indicates which color was presented. This rotation decomposes the input into two components: cos(*θ*) determines how much remains aligned with the color dimension, while sin(*θ*) *· s*_color_ determines how much projects onto the action dimension. By rotating in opposite directions (+*θ* vs -*θ*) for the two contexts, the action-aligned component reverses sign betweencontexts, directly implementing the context-dependent color-to-action mapping. For example, at *θ* = 0, inputs remain purely on the color axis; at *θ* = 90, inputs project entirely onto the action axis, eliminating color information. We swept *θ* between 0 and 90 and computed color and action variance from dPCA to assess how the rotation angle affects the representation of task variables.

#### Recurrent network model

We created a 4-area recurrent neural network model to solve the checkerboard task. We modeled activities of anterior, ventral, dorsal DLPFC and PMd activities. The state activity *x* of each unit is specified by the following equation:

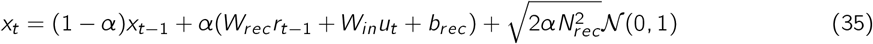

The firing rate of each recurrent unit is obtained by passing state activity through a relu nonlinearity:

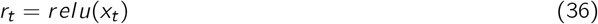

The output of the network *y*_*t*_ is given by:

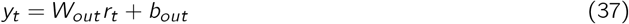

For each trial, we simulate *t* ∈ [0, 130) time points with 10 ms intervals giving us a total time of 1300 ms. Targets onset is at 100ms (t = 10), and checkerboard appear at 800ms (t = 80). Input *u*^130×3^ is three dimensional:

1st dimension: left target color (green: -1, red: 1).

2nd dimension: right target color (green: -1, red: 1).

3rd dimension: checkerboard color coherence *Coh* ∈ [−1, 1] (pure green: -1, pure red: 1)

First two dimensions of input turned on at target onset (*t* = 10) and third dimension of input turned on at checkerboard onset (*t* = 80).

The desired output *y* ^130×1^ is a one dimensional vector representing the chosen direction: -1 (left) and 1 (right) after checkerboard onset (*t >* 80). Before checkerboard, *y* = 0.

Each area contains 100 recurrent units. We applied 80/20 Dale’s law on connectivity matrix *W*_*rec*_ as first 80 units are excitatory and last 20 units are inhibitory. Based on anatomical evidences, we engineered inter-areal connections between areas. We enabled sparse feedforward connections from area 1 to area 2, area 2 to area 3 and area 3 to area 4. We enabled sparse feedback connections from area 2 to area 1, area 3 to area 1, area 3 to area 2 and area 4 to area 3. All feedforward and feedback connections among areas are excitatory to excitatory. We enabled input to area 1 and area 2 and output from only area 4 (Fig. S7a).

We modified this inter-areal connectivity motif to build 4 model frameworks: full model and 3 variants (Fig. S7b).

##### Full model

We created 3 separate compartments in area 1 and 2: input units (receives color and target configuration inputs), recurrent units and feedforward units. 30 excitatory units in area 1 and 10 excitatory units in area 2 receive color inputs. 10 excitatory units in area 1 and 30 excitatory units in area 2 receives target configuration (context). Note that 10 units in area 1 and 2 each receive both color and target configuration inputs. 10 excitatory units in area 1 and area 2 apart from 30 input units are recurrent units. The rest 40 excitatory units sparsely feedforward to random excitatory units in next area. All feedback connections are random excitatory units to random excitatory units (Fig. S7a).

**Variant 1:** Same as full model but we removed all feedback connections among areas.

**Variant 2:** Same as full model but there is no target configuration input to area 1.

**Variant 3:** There is no compartments in area 1 and 2. Units receiving color input, configuration input and feedforward units are totally random and can overlap.

We used back propagation through time with a mean squared error loss function between real output and desired output y to train the networks. We evaluate the final decision of the model as the 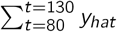 after checkerboard onset. The decision is left if 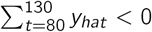 and right if 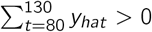. We trained 30 models for each framework. Each training set contains 8000 trials; each testing set contains 2000 trials.

For each trained model, we normalized each unit’s *r*_*t*_ using same method described in Methods d and performed dPCA on all 400 units together. For each model framework, we combined 30 trained models per area (100×30) and calculated the mean dPC loadings and SEM. To quantify color and color x action variance explained in each area, we performed dPCA on each area separately for all of the 30 trained models.

## Acknowledgments

We thank the Whitehall Foundation (2019-12-77), the Brain and Behavior Research Foundation (27923), the National Institute of Neurological Disorders and Stroke (R00 NS092972, R01 NS121409, R01 NS122969, R01 NS135361, F31 NS131018), and the Moorman Simon Interdisciplinary Career Fellowhsip fund at Boston University for enabling this work. We thank Nabil Abu Nassar (NAN Instruments) for providing us with custom drives for Neuropixels Recordings.

## References

[1] Chandrasekaran C, Peixoto D, Newsome WT, Shenoy KV (2017) Laminar differences in decision-related neural activity in dorsal premotor cortex. Nature Communications 8(1):614.

[2] Boucher PO, et al. (2023) Initial conditions combine with sensory evidence to induce decision-related dynamics in premotor cortex. Nature Communications 14(1):6510.

[3] Bennur S, Gold JI (2011) Distinct representations of a perceptual decision and the associated oculomotor plan in the monkey lateral intraparietal area. The Journal of Neuroscience: The Official Journal of the Society for Neuroscience 31(3):913–921.

[4] Charlton JA, Goris RL (2024) Abstract deliberation by visuomotor neurons in prefrontal cortex. Nature Neuroscience 27(6):1167–1175.

[5] Kira S, Safaai H, Morcos AS, Panzeri S, Harvey CD (2023) A distributed and efficient population code of mixed selectivity neurons for flexible navigation decisions. Nature communications 14(1):2121.

[6] Rigotti M, et al. (2013) The importance of mixed selectivity in complex cognitive tasks. Nature 497(7451):585–590.

[7] Gold JI, Shadlen MN (2007) The neural basis of decision making. Annual Review of Neuroscience 30(1):535–574.

[8] Shadlen MN, Kiani R (2013) Decision making as a window on cognition. Neuron 80(3):791–806.

[9] Brody CD, Hanks TD (2016) Neural underpinnings of the evidence accumulator. Current Opinion in Neurobiology 37:149–157.

[10] Hanks TD, et al. (2015) Distinct relationships of parietal and prefrontal cortices to evidence accumulation. Nature 520(7546):220–223.

[11] Steinmetz NA, Zatka-Haas P, Carandini M, Harris KD (2019) Distributed coding of choice, action and engagement across the mouse brain. Nature 576(7786):266–273.

[12] Okazawa G, Kiani R (2023) Neural Mechanisms That Make Perceptual Decisions Flexible. Annual Review of Physiology 85:191–215.

[13] Rosen MC, Freedman DJ (2025) How distributed is the brain-wide network that is recruited for cognition? Nature Reviews Neuroscience pp. 1–13.

[14] Gupta D, et al. (2024) A multi-region recurrent circuit for evidence accumulation in rats. BioRxiv.

[15] Steinmetz NA, Zatka-Haas P, Carandini M, Harris KD (2019) Distributed coding of choice, action and engagement across the mouse brain. Nature pp. 1–8.

[16] Roitman JD, Shadlen MN (2002) Response of neurons in the lateral intraparietal area during a combined visual discrimination reaction time task. The Journal of Neuroscience 22(21):9475–9489.

[17] Cisek P, Kalaska JF (2005) Neural correlates of reaching decisions in dorsal premotor cortex: Specification of multiple direction choices and final selection of action. Neuron 45(5):801–814.

[18] Badre D, D’esposito M (2009) Is the rostro-caudal axis of the frontal lobe hierarchical? Nature reviews neuroscience 10(9):659–669.

[19] Basile C, et al. (2025) Encoding of visual stimuli and behavioral goals in distinct anatomical areas of monkey ventrolateral prefrontal cortex. PLoS Biology 23(8):e3003041.

[20] Freedman DJ, Assad JA (2011) A proposed common neural mechanism for categorization and perceptual decisions. Nature Neuroscience 14(2):143–146. Number: 2 Publisher: Nature Publishing Group.

[21] Sandhaeger F, Omejc N, Pape AA, Siegel M (2023) Abstract perceptual choice signals during action-linked decisions in the human brain. Plos Biology 21(10):e3002324.

[22] Steinmetz NA, Moore T (2014) Eye movement preparation modulates neuronal responses in area v4 when dissociated from attentional demands. Neuron 83(2):496–506.

[23] Miller EK, Cohen JD (2001) An Integrative Theory of Prefrontal Cortex Function. Annual Review of Neuroscience 24(1):167–202. _eprint:.

[24] Siegel M, Buschman TJ, Miller EK (2015) Cortical information flow during flexible sensorimotor decisions. Science (New York, N.Y.) 348(6241):1352–1355.

[25] Thura D, Cisek P (2014) Deliberation and commitment in the premotor and primary motor cortex during dynamic decision making. Neuron 81(6):1401–1416.

[26] Tye KM, et al. (2024) Mixed selectivity: Cellular computations for complexity. Neuron 112(14):2289– 2303.

[27] Dang W, Jaffe RJ, Qi XL, Constantinidis C (2021) Emergence of nonlinear mixed selectivity in prefrontal cortex after training. Journal of Neuroscience 41(35):7420–7434.

[28] Mendoza-Halliday D, et al. (2024) A ubiquitous spectrolaminar motif of local field potential power across the primate cortex. Nature Neuroscience 27(3):547–560.

[29] Fu Z, et al. (2022) The geometry of domain-general performance monitoring in the human medial frontal cortex. Science 376(6593):eabm9922.

[30] Courellis HS, et al. (2024) Abstract representations emerge in human hippocampal neurons during inference. Nature 632(8026):841–849.

[31] Fusi S, Miller EK, Rigotti M (2016) Why neurons mix: high dimensionality for higher cognition. Current opinion in neurobiology 37:66–74.

[32] Kobak D, et al. (2016) Demixed principal component analysis of neural population data. eLife 5.

[33] Bichot NP, Heard MT, DeGennaro EM, Desimone R (2015) A source for feature-based attention in the prefrontal cortex. Neuron 88(4):832–844.

[34] Gerbella M, Borra E, Tonelli S, Rozzi S, Luppino G (2013) Connectional heterogeneity of the ventral part of the macaque area 46. Cerebral Cortex 23(4):967–987.

[35] Pagan M, et al. (2025) Individual variability of neural computations underlying flexible decisions. Nature 639(8054):421–429.

[36] Soldado-Magraner J, Mante V, Sahani M (2024) Inferring context-dependent computations through linear approximations of prefrontal cortex dynamics. Science Advances 10(51):eadl4743.

[37] Murray JD, et al. (2017) Stable population coding for working memory coexists with heterogeneous neural dynamics in prefrontal cortex. Proceedings of the National Academy of Sciences 114(2):394–399.

[38] Siegel M, Buschman TJ, Miller EK (2015) Cortical information flow during flexible sensorimotor decisions. Science 348(6241):1352–1355.

[39] Kleinman M, et al. (2025) The information bottleneck as a principle underlying multi-area cortical representations during decision-making. eLife.

[40] Mante V, Sussillo D, Shenoy KV, Newsome WT (2013) Context-dependent computation by recurrent dynamics in prefrontal cortex. Nature 503(7474):78–84.

[41] Romanski LM (2007) Representation and integration of auditory and visual stimuli in the primate ventral lateral prefrontal cortex. Cerebral Cortex 17(Suppl_1):i61–i69.

[42] Hoshi E (2006) Functional specialization within the dorsolateral prefrontal cortex: a review of anatomical and physiological studies of non-human primates. Neuroscience research 54(2):73–84.

[43] Wilson FA, Scalaidhe SP, Goldman-Rakic PS (1993) Dissociation of object and spatial processing domains in primate prefrontal cortex. Science 260(5116):1955–1958.

[44] Gou L, et al. (2025) Single-neuron projectomes of macaque prefrontal cortex reveal refined axon targeting and arborization. Cell 188(14):3806–3822.

[45] Chandrasekaran C, et al. (2025) No central executive? decision formation through multi-area population dynamics. Journal of Neuroscience 45(46).

[46] Thura D, Cabana JF, Feghaly A, Cisek P (2022) Integrated neural dynamics of sensorimotor decisions and actions. PLOS Biology 20(12):e3001861.

[47] Toth LJ, Assad JA (2002) Dynamic coding of behaviourally relevant stimuli in parietal cortex. Nature 415(6868):165–168.

[48] Freedman DJ, Assad JA (2009) Distinct encoding of spatial and nonspatial visual information in parietal cortex. Journal of Neuroscience 29(17):5671–5680.

[49] Okazawa G, Kiani R (2025) Inferotemporal cortex encodes formation and termination of perceptual decisions. bioRxiv pp. 2025–12.

[50] Tian LY, et al. (2025) Neural representation of action symbols in primate frontal cortex. bioRxiv.

[51] Javadzadeh M, Schimel M, Hofer SB, Ahmadian Y, Hennequin G (2024) Dynamic consensus-building between neocortical areas via long-range connections. bioRxiv pp. 2024–11.

[52] Galgali AR, Sahani M, Mante V (2023) Residual dynamics resolves recurrent contributions to neural computation. Nature Neuroscience 26(2):326–338.

[53] Chandrasekaran C, Bray XE, Shenoy KV (2019) Frequency shifts and depth dependence of premotor beta band activity during perceptual decision-making. Journal of Neuroscience 39(8):1420–1435.

[54] Trautmann EM, et al. (2019) Accurate Estimation of Neural Population Dynamics without Spike Sorting. Neuron.

