## Supplementary materials for "Distinct neural dynamics in prefrontal and premotor cortex during flexible perceptual decisions"

### Supplementary Information for *Distinct neural dynamics in prefrontal and premotor cortex during flexible perceptual decisions*

Tian Wang[a], Eric Kenji Lee[b], Nicole Carr[a], Yuke Li[a], Chandramouli Chandrasekaran[a,b,c,d,*]

[a] *Department of Biomedical Engineering, Boston University, Boston, 02115, MA, USA*
[b] *Department of Psychological and Brain Sciences, Boston University, Boston, 02115, MA, USA*
[c] *Center for Systems Neuroscience, Boston University, Boston, 02115, MA, USA*
[d] *Department of Anatomy & Neurobiology, Boston University, Boston, 02118, MA, USA*

**Contents**

**Appendix A: Nonlinear Mixed Selectivity** S-2

**Fig. S1 - Localization of DLPFC recording sites** S-4

**Fig. S2 - Selectivity of single neurons** S-5

**Fig. S3 - Mixed selectivity in other PC dimensions** S-6

**Fig. S4 - Types of mixed selective and pure selective representations and predictions** S-7

**Fig. S5 - Functional gradients** S-8

**Fig. S6 - Inputs and dynamics for different models in Fig. 6** S-9

**Fig. S7 - A 4-area recurrent neural network model can explain the gradient** S-10

#### Appendix A: Nonlinear-Mixed Selectivity

One strategy for solving the XOR task is to generate high-dimensional neural representations that unmix color and action. Such high-dimensional representations necessitate that a brain area encodes multiple relevant variables. This could take three forms that can be naturally quantified in a two-way ANOVA (Table S1).

Neurons could be purely tuned for color choice (**color**, top row in Table S1) or purely tuned for action choice (**action**, second row in Table S1) or a linear combination of both effects, a phenomenon termed linear mixed selectivity.

| Selectivity Type | Color | Action | Interaction |
|---|---|---|---|
| Classical (CS) | ✓ | – | – |
| | – | ✓ | – |
| Linear Mixed (LMS) | ✓ | ✓ | – |
| Target Configuration (Pattern 1 below) (NMS) | – | – | ✓ |
| Nonlinear Mixed (Pattern 2-4 below) (NMS) | ✓ | ✓ | ✓ |

Table S1: Classification of neuronal selectivity based on two-way ANOVA

Neurons could also be sensitive to nonlinear mixtures of color and action, and thus have nonlinear mixed selectivity[26]. In a two-way ANOVA the interaction term tests whether the effect of color depends on action (or vice versa). A significant interaction term therefore indicates nonlinear relationships, but can arise from either encoding the target configuration or a more sophisticated pattern (Table S1, bottom two rows). The section below expands on how such nonlinear mixed selectivity could occur in our task.

*Interaction terms in a 2-way ANOVA*

The interaction sum of squares is:

$$SS_{interaction} = n \sum_{i,j} (\bar{Y}_{ij} - \bar{Y}_{i.} - \bar{Y}_{.j} + \bar{Y}_{..})^2 \quad (.1)$$

where $Y$ represents the firing rate of a neuron at a particular time point, $\bar{Y}_{ij}$ is the mean firing rate for color $i$ and action $j$, $\bar{Y}_{i.}$ and $\bar{Y}_{.j}$ are the marginal means, $\bar{Y}_{..}$ is the grand mean, and $n$ is the number of trials per condition.

For a 2×2 design with color choice and action choice, this simplifies to:

$$SS_{interaction} = \frac{n}{4}[(\bar{Y}_{RL} - \bar{Y}_{RR} - \bar{Y}_{GL} + \bar{Y}_{GR})^2] \quad (.2)$$

The interaction is significant when responses to the four color-action combinations cannot be predicted by simply adding the main effects of color and action.

*Examples of Interaction Patterns*

*Pattern 1: Selectivity to Target Configuration*

Neurons that are sensitive to the target configuration would show significant interaction terms and are examples of neurons that could help solve flexible decisions. For instance, Table S2 shows an example of a neuron that would have no effect of color and choice but would have a significant interaction term.

| | Left | Right |
|---|---|---|
| Red | **20** | 5 |
| Green | 5 | **20** |

Table S2: Example of a target configuration sensitive neuron

The marginal means for color and choice are equal (12.5) and thus there are no main effects of action or color choice. In contrast, the interaction term is $(20-5-5+20)^2 = 900$, which is consistent with encoding target configuration. Such neurons contribute to solving the XOR computation by encoding which context the animal is in.

*Pattern 2: tuning to a specific combination of color choice and action choice*
Another example of a significant interaction is a neuron that prefers a particular combination of color and action. For instance, a neuron that fires strongly only for the red-left case (Table S3).

| | Left | Right |
|---|---|---|
| Red | **25** | 5 |
| Green | 5 | 5 |

Table S3: Example of a neuron selective for red-left

The marginal means are: Red = 15, Green = 5, Left = 15, Right = 5. Such a neuron would have main effects for both color choice and action choice. The interaction term is $(25 - 5 - 5 + 5)^2 = 400$. This neuron is mixed-selective and could contribute to the representations found in Fig. 4a.

*Pattern 3: tuning to 3 specific combination of color choice and action choice*
An example of a significant interaction is a neuron that can differentiate 3 particular combinations of color and action, shown in (Table S4).

| | Left | Right |
|---|---|---|
| Red | **25** | 10 |
| Green | 10 | 5 |

Table S4: Example of a neuron that differentiates 3 color/action combinations

The marginal means are: Red = 17.5, Green = 7.5, Left = 17.5, Right = 7.5. Such a neuron would have main effects for both color choice and action choice. The interaction term is $(25 - 10 - 10 + 5)^2 = 100$. This neuron is mixed-selective and could contribute to the representations found in Fig. 4a.

*Pattern 4: each color choice and action choice has different tunings*
An example of a significant interaction is a neuron that differentiates any particular combination of color and action, shown in (Table S5).

| | Left | Right |
|---|---|---|
| Red | **25** | 20 |
| Green | 15 | 5 |

Table S5: Example of a neuron selective to all color/action combinations

The marginal means are: Red = 22.5, Green = 20, Left = 20, Right = 12.5. Such a neuron would have main effects for both color choice and action choice. The interaction term is $(25 - 20 - 15 + 5)^2 = 25$. This neuron is mixed-selective and could contribute to the representations found in Fig. 4a.

#### Supplementary Figures

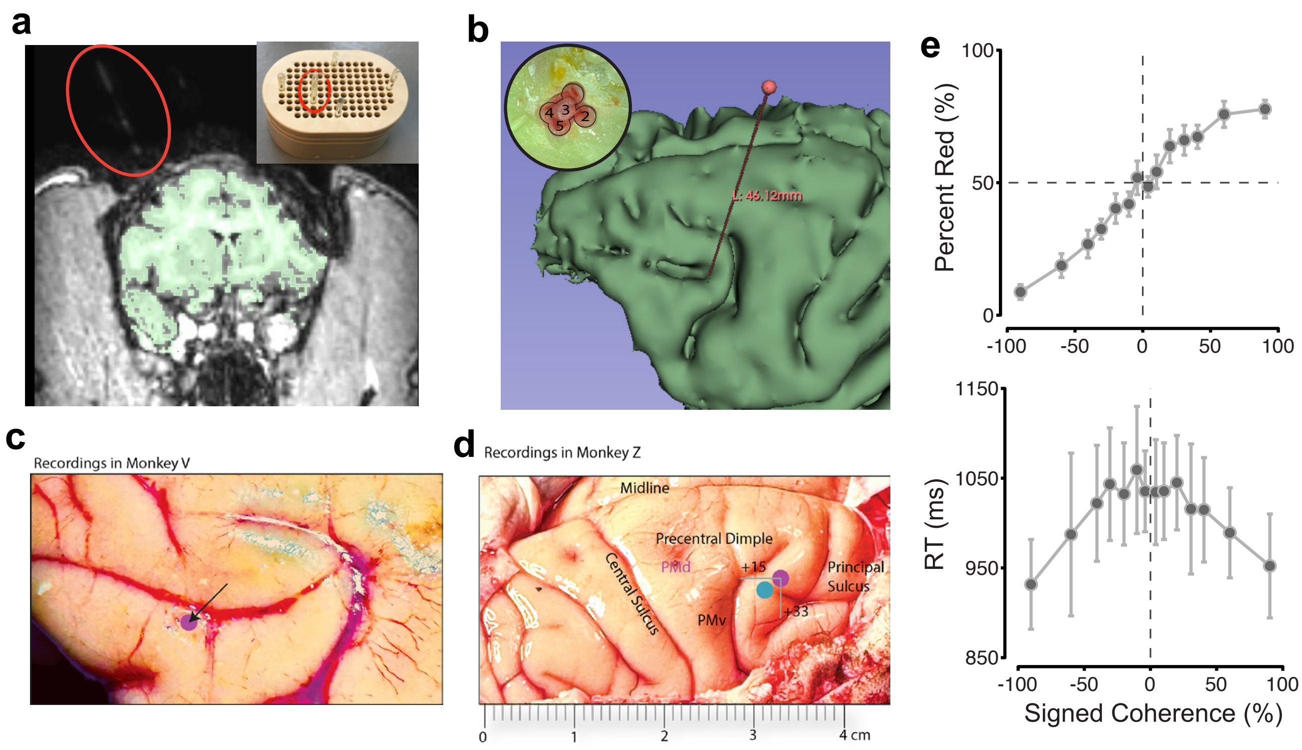


Figure S1: **Localization of DLPFC recording sites in monkeys T, V, and Z using MRI and histology** (**a**) A coronal view of MRI for monkey T. Green area is the brain segmented from a T1 MRI scan. Inset on right top corner is a vitamin E capillary tube (red circle) placed in the grid. We placed the grid on monkey's chamber to localize the recording site. (**b**) Segmented brain from a T1 scan and localization of recording site in monkey T. Inset on left top corner is the burrhole number. According to MRI localization and grid location of each day's recording, we estimated posterior edge of burrhole 3 as posterior end of principal sulcus. (**c-d**) Monkey V and Z brain image with recording sites. Recording sites are marked with purple and blue dots. (**e**) Monkey Z psychometric curve and reaction curve. We only incorporate monkey Z's data in Fig. 5.

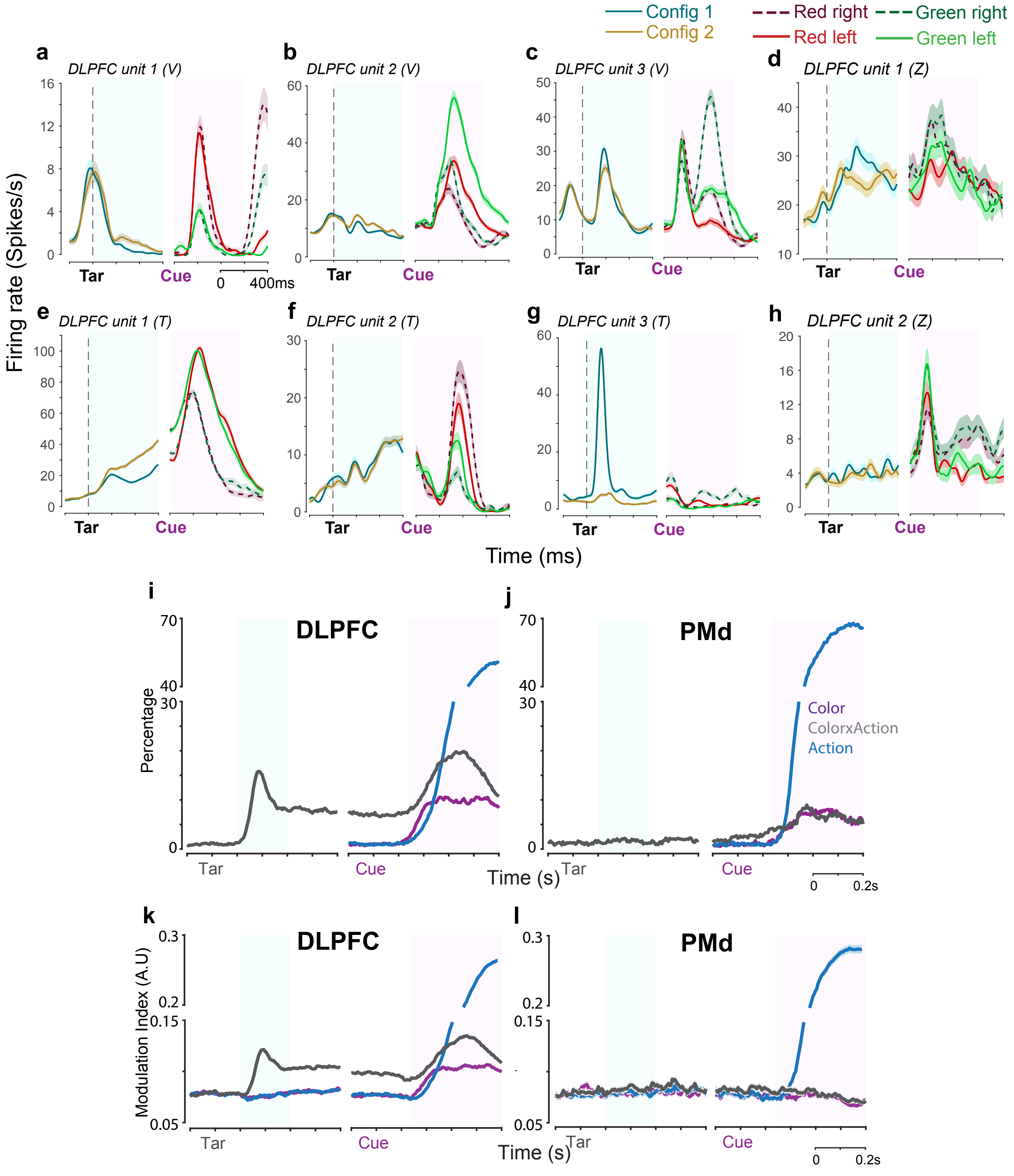


Figure S2: **Mixed selectivity in DLPFC units and temporal evolution of task variable encoding** (**a-h**) Additional examples of DLPFC units demonstrating diverse patterns of mixed selectivity necessary for solving the XOR task. Shaded regions indicate SEM. Vertical dashed lines mark target onset (Tar). (**i-j**) Time-resolved percentage of significantly modulated units in DLPFC (g) and PMd (h) from a two-way ANOVA (color and action as main effects, $p < 0.01$) for each neuron using a sliding window (50 ms window, 5 ms step). In DLPFC, color (purple), choice (blue), and interaction term (NMS, color $\times$ action, gray) modulation are all robust. When main effects are absent (before Cue), the interaction between color $\times$ action is essentially target configuration. In PMd, modulation during the target epoch is minimal. Action modulation dominates after checkerboard onset, reaching 70% of units, while color and interaction modulation are modest throughout the trial. (**k-l**) Average effect size of task variable modulation over time in DLPFC (i) and PMd (j) for all neurons. Modulation index quantifies the normalized difference in firing rates between conditions (e.g. for color, $(FR_{red} - FR_{green})/(FR_{red} + FR_{green})$). In DLPFC, all task variables show substantial effect sizes that build after their respective onsets. In PMd, only action exhibits large effects (>0.2), while color, configuration, and interaction modulation indices remain low.

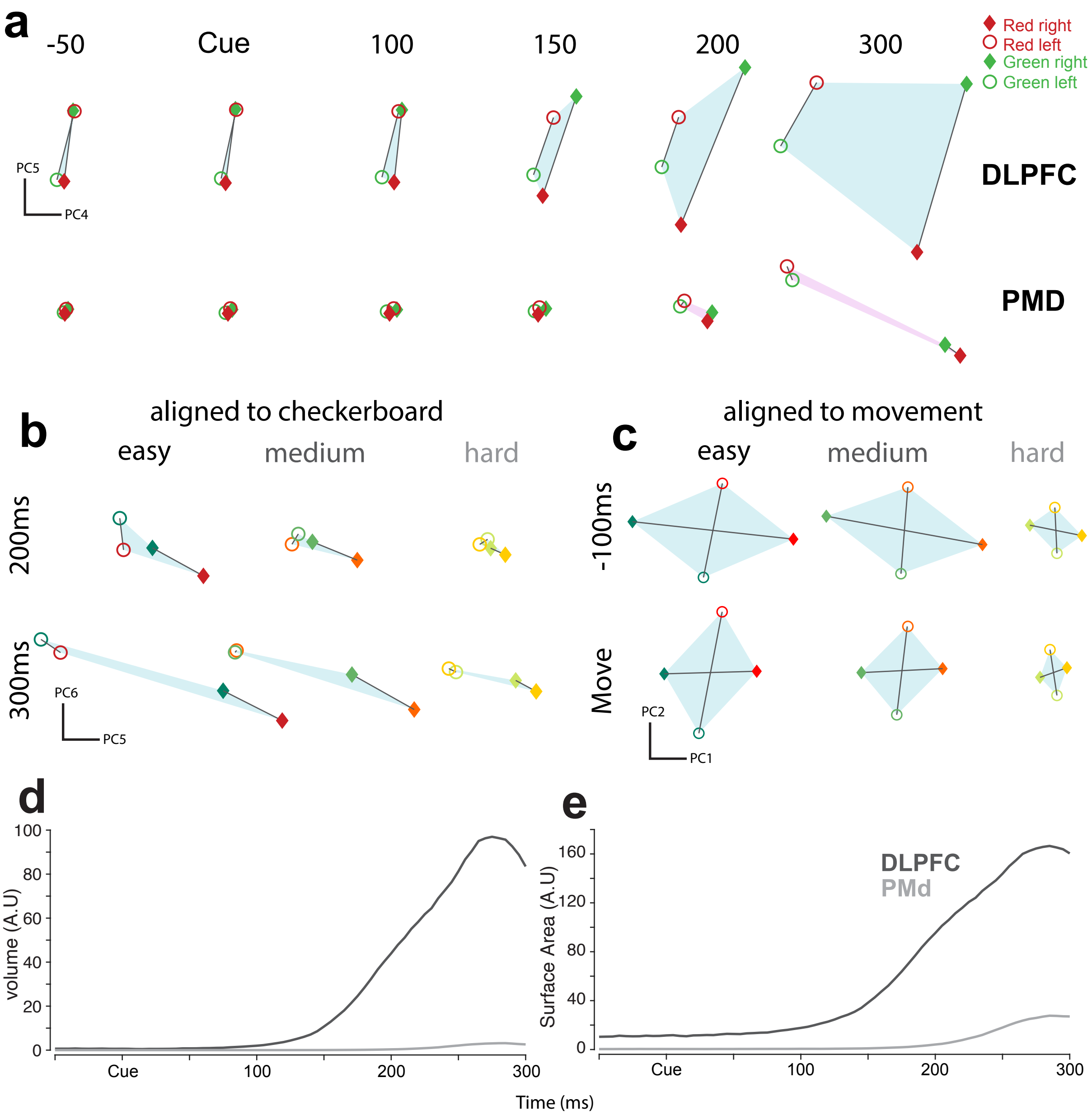


Figure S3: **Mixed selective high-dimensional representations in DLPFC and PMd** (**a**) Firing rates projected on to PC4 and PC5 for DLPFC (top) and PMd (bottom) at different time points around checkerboard onset. Same as Fig. 4a, four color/action combinations are well separated in DLPFC. The quadrilateral shape shows linear action choice separation with no pure color separation. In PMd, only action can be well separated. (**b**) DLPFC population dynamics projected to PC5 and PC6 for easy, medium and hard trials at 200 ms (top row) and 300 ms (bottom row) after checkerboard onset. Action choice is the major signal in these PC projections. The distance between left and right choices, representing the amplitude of action choice, decreases from easy to hard difficulty. (**c**) DLPFC population dynamics projected to PC1 and PC2 (after action signal removal) at -100 ms and 0 ms aligns to movement onset. Same as Fig. 4c, quadrilateral shape also decreases as task difficulty increases, representing mixed selectivity signal is not only delayed but also weakens in more difficult trials. (**d**) The time resolved volume of the tetrahedron embedded by top 10 PC space bounded by 4 color/action combinations. The tetrahedron volume quantifies nonlinear mixed selectivity of color and action. In DLPFC, tetrahedron volume increases rapidly start from 100 ms after checkerboard onset and peaks before 300 ms. PMd tetrahedron volume does not increase. (**e**) The time resolved surface area of the tetrahedron bounded by 4 color/action combinations embedded in the subspace of the top 10 PCs space . Similar to volume, the tetrahedron surface area increases rapidly in DLPFC but not in PMd.

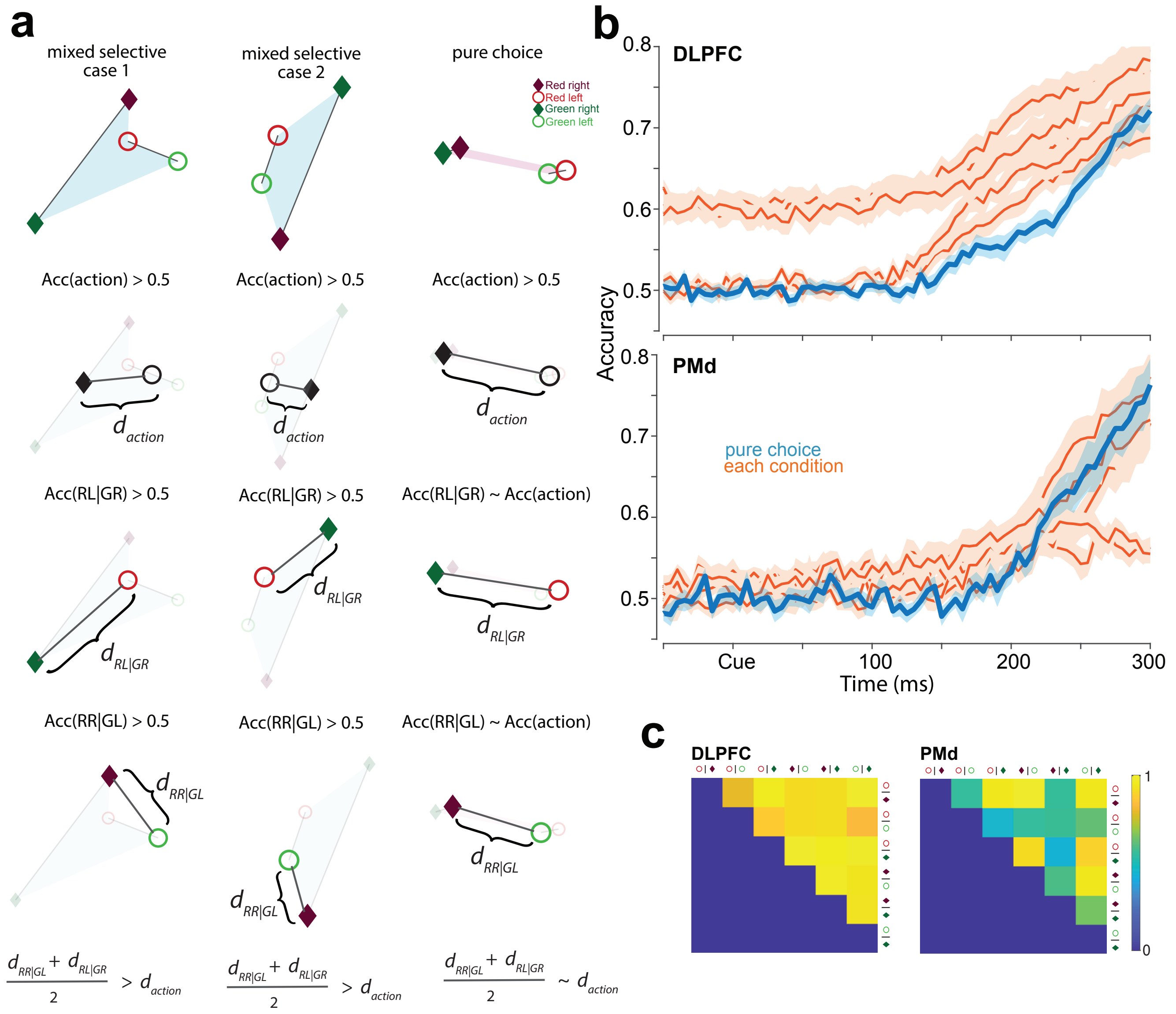


Figure S4: **Types of mixed selective representations** (**a**) Schematic illustrating how nonlinear choice decoding can exceed pure choice decoding. Top row shows representational geometry for three scenarios: mixed selective case 1 (left, with linear color separation), mixed selective case 2 (middle, without linear color separation), and pure choice (right). In all cases, pure choice decoding accuracy (Acc(action) > 0.5, second row) distinguishes left versus right choices by comparing average positions of RL, GL versus RR, GR (distance $d_{\text{action}}$). Nonlinear choice decoding (third and fourth rows) compares correct versus incorrect outcomes: RL, GR versus RR, GL, quantified as the average of $d_{\text{RL|GR}}$ and $d_{\text{RR|GL}}$. When all four conditions are well-separated (left and middle columns), these pairwise distances exceed $d_{\text{action}}$, indicating nonlinear mixed selectivity. In pure choice representations (right), pairwise distances equal $d_{\text{action}}$. (**b**) Logistic regression decoding accuracy for all six pairwise comparisons between the four color-action combinations. In DLPFC (top), all pairwise accuracies increase after checkerboard onset and exceed pure choice decoding, with nonlinear choice pairs (Acc(RL|GR) and Acc(RR|GL), orange) exceeding action-only decoding (blue). This indicates separation of all four conditions, consistent with mixed selective representations. In PMd (bottom), four pairwise comparisons track pure choice decoding, while two pairs show minimal increases, indicating primarily action-only representations. Shaded regions show SEM across sessions. (**c**) Correlation matrices showing $R^2$ from linear regression comparing session-averaged decoding accuracy between each pair of the six comparisons. In DLPFC, high $R^2$ values across all pairs indicate that all pairwise decoding accuracies increase together, reflecting consistent separation of all four conditions. In PMd, lower $R^2$ values between some pairs indicate heterogeneous decoding patterns, with some pairs tracking action-only representations. These results demonstrate stronger mixed selective representations in DLPFC compared to PMd.

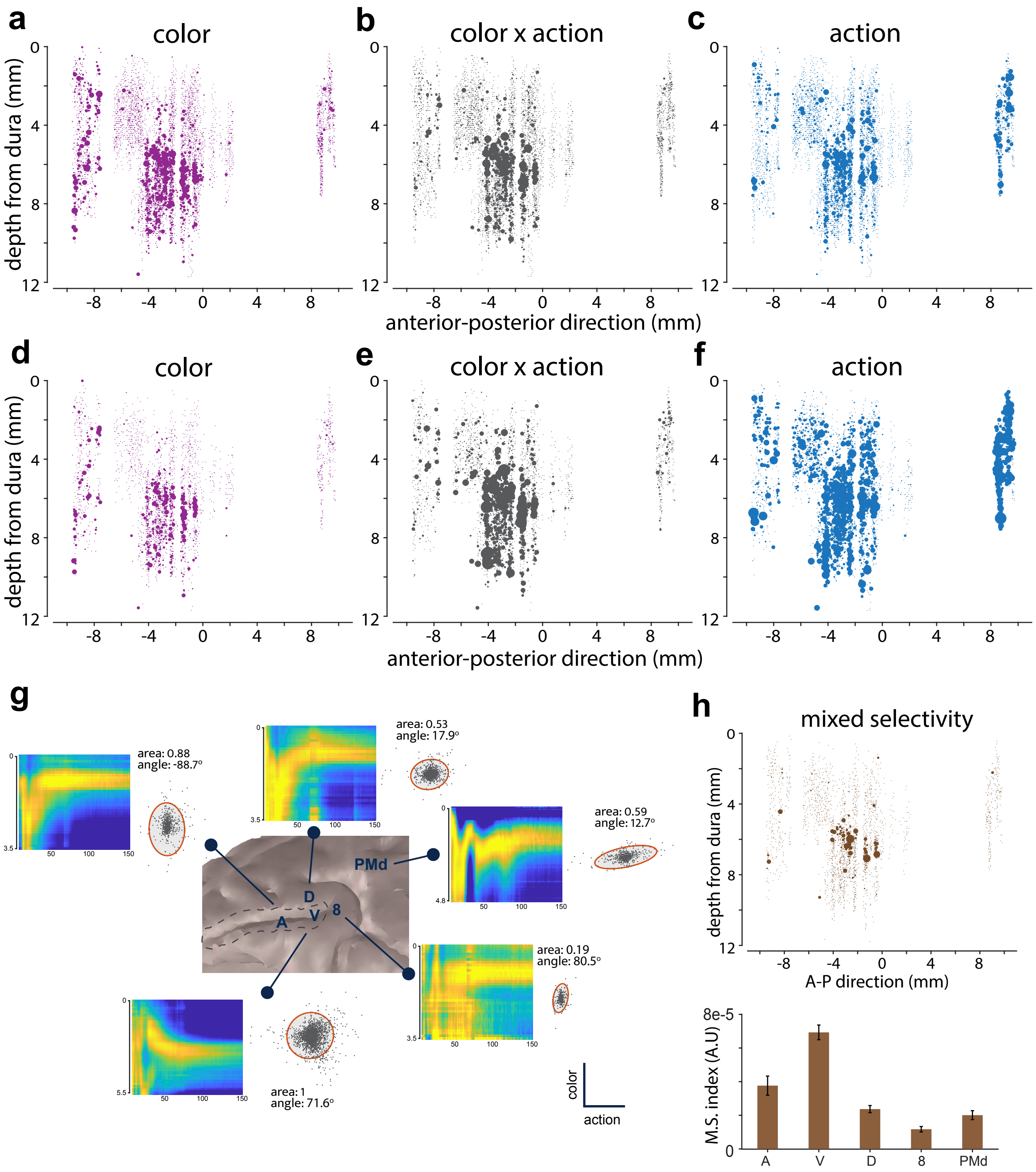


Figure S5: **Functional gradients in DLPFC and PMd** (**a-c**) dPC loading of each unit similar to Fig. 5a-c. The only difference is that we performed dPCA without normalizing the binned firing counts of each unit. (**d-f**) Each unit's effect size reported as average Cohen's d across a time window [150,400] after checkerboard onset. (**g**) Average spectrolaminar motif of each recorded area (same as Fig. 5d) and scatter plot of dPC loadings on action and color axis. The ellipse covers 99% of the covariance. We calculated the angle and area of each ellipse, normalized relative to ventral DLPFC. Anterior DLPFC shows the strongest color signals, while PMd is most selective for action. (**h**) Top: the scatter plot of each unit's mixed selectivity index (defined in *Methods: Anatomical localization of recording sites*). Dot size quantifies how mixed selective a unit is. Bottom: bar graph shows each areas average mixed selectivity index (error bar is SEM).

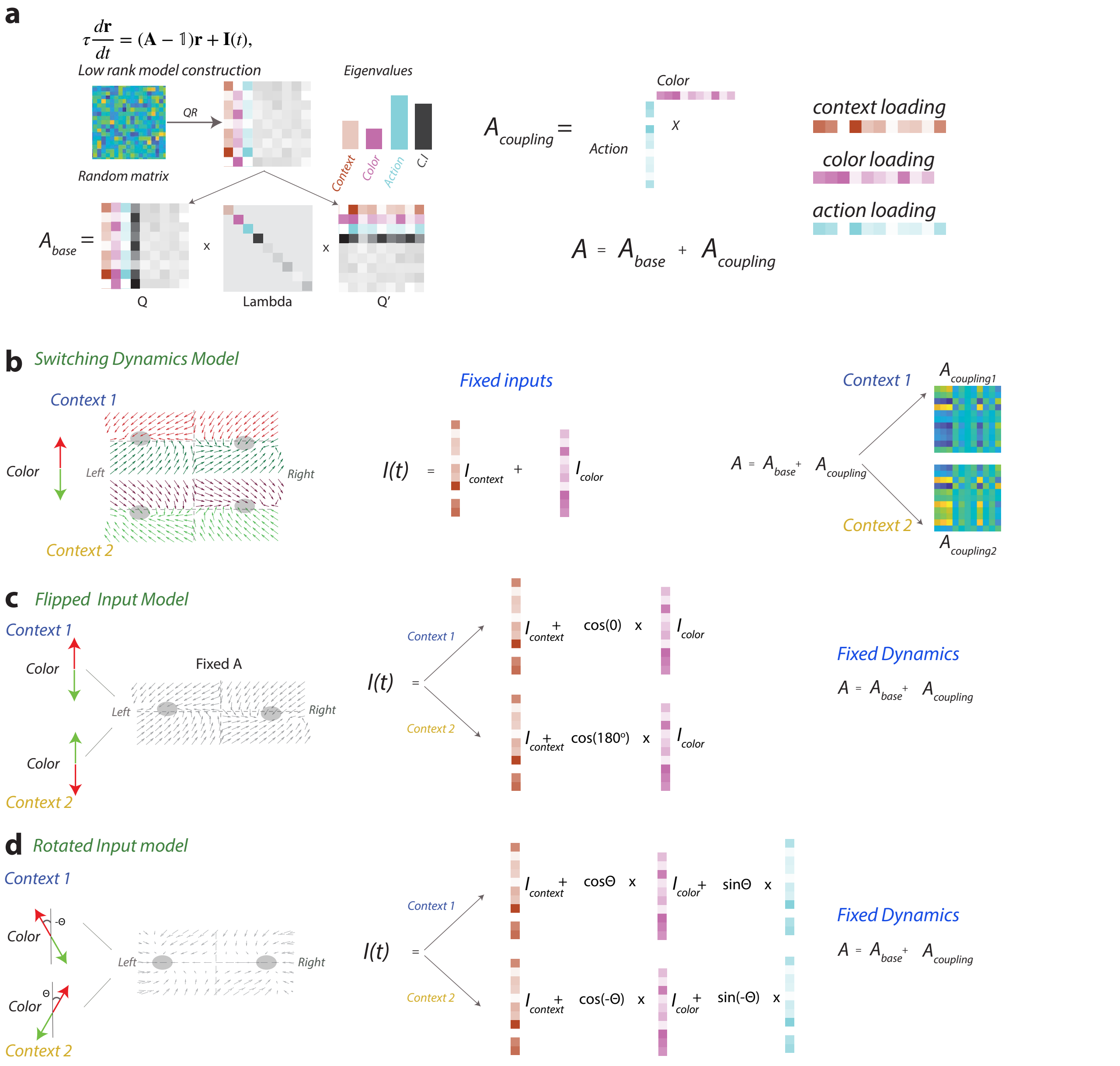


$$\tau \frac{d\mathbf{r}}{dt} = (\mathbf{A} - \mathbb{1})\mathbf{r} + \mathbf{I}(t),$$

Figure S6: **Schematic of model construction and variants of low-rank models** (**a**) The connectivity matrix **A** consists of two components: $\mathbf{A}_{\text{base}}$ and $\mathbf{A}_{\text{coupling}}$. $\mathbf{A}_{\text{base}} = \mathbf{Q}\mathbf{\Lambda}\mathbf{Q}^T$, where **Q** is an orthonormal basis defining four computational dimensions: context, color, choice, and condition-independent signal. $\mathbf{\Lambda}$ specifies eigenvalues that determine the temporal stability of each dimension. $\mathbf{A}_{\text{coupling}}$ is a rank-1 matrix that couples color and choice dimensions. (**b**) *Switching dynamics model*: Inputs are fixed across contexts and align with their corresponding dimensions in **Q**, with heterogeneous tuning weights for each neuron. Context modulates the computation by reversing the sign of $\mathbf{A}_{\text{coupling}}$, switching between $\mathbf{A}_1$ and $\mathbf{A}_2$. (**c**) *Flipped input model*: The connectivity matrix **A** is fixed. Context modulates the computation by inverting the sign of the color input (180° flip), effectively reversing the color-to- action mapping while preserving heterogeneous tuning. (**d**) *Rotated input model*: The connectivity matrix **A** is fixed. Context modulates the computation by rotating the color input symmetrically by $\pm\theta$ in the color-choice plane. This creates context-dependent projections onto both color and choice dimensions, with the choice-aligned component reversing sign between contexts to implement the XOR mapping.

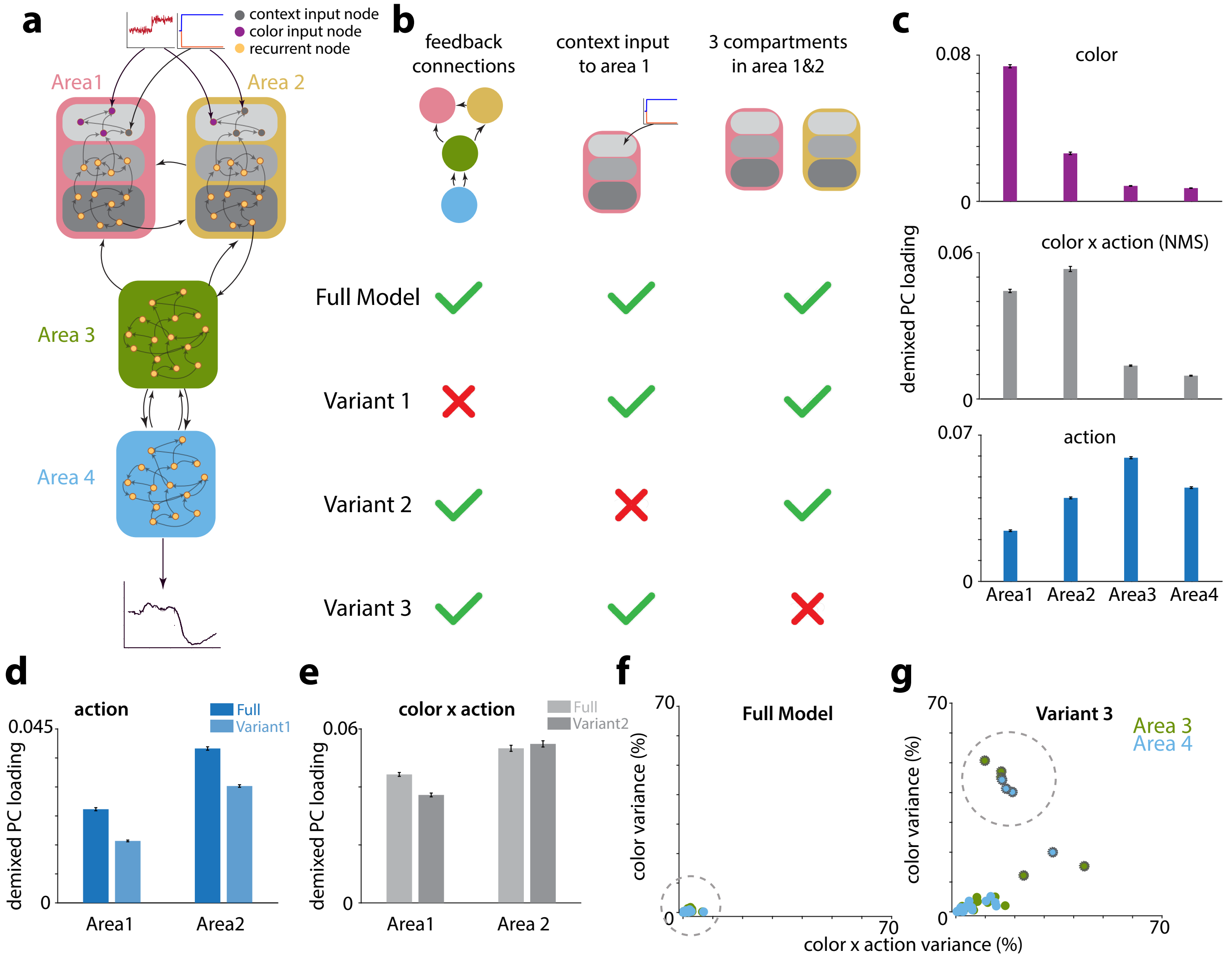


Figure S7: **4-Area recurrent neural network model** (**a**) Full 4-area RNN model structure. Area 1 and 2 has 3 compartments: input units, recurrent units, feedforward units. 30 units in area 1 and 10 units in area 2 receives color input; 10 units in area 1 and 30 units in area 2 receive context inputs. 40 feedforward units project to next area. Feedback is from random units of current area to random units in previous area. (**b**) 3 variants of RNN model structure: variant 1: no feedback connections; variant 2: no context input to area 1; variant 3: no compartments in area 1 and 2 (**c**) Average dPC loading of each color (top), color × action (middle) and action (bottom) across all 30 sessions of the full model. Error bar is SEM. Color dPC loadings decreases from area 1 to 4; area 2 has highest color and choice mixed selectivity loadings. RNN model also shows a functional gradient similar to neural data. (**d**) Comparison of average action dPC loadings in area 1 and 2 between model and variant 1 (no feedback). On average, choice representation is weaker with no feedback connections. (**e**) Comparison of average color × action dPC loadings in area 1 and 2 between model and variant 2 (no context input in area 1). On average, area 2 nonlinear mixed selectivity increases if area 1 receives no context inputs. (**f-g**) Comparison of color × action and color variance explained in area 3 and 4 between model and variant 3 (no compartments). We trained 30 models for vanilla model and variant 3; each dot is one trained model. Feedforward units segregating from input units ensures minimal input signal representation in area 3 and 4, while some trained models in variant 3 have high color and color × action signal (dots with gray rims).
